# Melatonin-mediated methylglyoxal homeostasis and regulation of autophagy during seed germination under PEG-induced drought stress in upland cotton

**DOI:** 10.1101/2025.02.18.638685

**Authors:** Deepika Dake, Laha Supriya, Amarjeet Kumar, Padmaja Gudipalli

## Abstract

Methylglyoxal (MGO), toxic byproduct of glycolysis, acts as a signaling molecule at low levels, but its over-accumulation during drought stress disrupts redox balance and accelerates cell death. Contrarily, melatonin maintains redox balance, particularly during stress. The redox status and MGO level might differ in drought-sensitive and drought-tolerant varieties, so shall the melatonin’s effect. The study evaluated the effect of melatonin-priming on MGO detoxification and autophagy during polyethylene glycol (PEG)-induced drought stress during seed germination in drought-sensitive (L-799) and drought-tolerant (Suraj) varieties of upland cotton. Melatonin-priming increased endogenous melatonin content, reduced MGO accumulation and advanced glycation end-products (AGEs), and downregulated the expression of MGO biosynthesis genes in L-799 under stress. The expression and activities of glyoxalases and non-glyoxalases were upregulated, showing melatonin’s effectiveness in MGO detoxification. Additionally, priming upregulated the expression of *TPI1*, *PGK5*, *and PK1* and downregulated *HK3* expression, allowing better conversion of glucose to pyruvate, leading to reduced MGO in L-799. The downregulation of necrosis-related genes with reduced cell death in L-799 shows the potential of priming in maintaining cell viability under stress. Furthermore, upregulated expression of *SnRK1.1*, *SnRK2.6* genes and KIN10 protein levels, with enhanced autophagy markers (*ATGs*, MDC-stained bodies, lipidated-ATG8), confirmed improved autophagy in melatonin-primed L-799 under stress. Despite lowered ABA, melatonin-mediated MGO homeostasis likely activated MAPK6, inducing autophagy independent of ABA in stressed plants. Conversely, Suraj, with higher endogenous melatonin and inherent stress tolerance, showed limited response to priming. Thus, the study illustrates melatonin’s role in regulating MGO homeostasis and autophagy under drought stress in cotton.

## 1. Introduction

Seed germination is a pivotal phase of plant life cycle that signifies the transition of a quiescent seed into a metabolically active state leading to the emergence of seedlings (El-Maarouf-Bouteau 2022). A healthy germination process ensures production of uniform and robust seedlings through efficient resource utilization and increasing photosynthetic potential that results in higher yields (Ali and Elozeiri 2017). Drought is a major abiotic stress factor that adversely affects seed germination process, and consequently diminishes seedling growth and their establishment (Martínez-Ballesta et al. 2020; Maksimović et al. 2021). Water unavailability disrupts the fundamental imbibition process during germination, impeding the enzymatic systems essential for nutrient mobilization (Bove et al. 2002). Additionally, drought stress causes intricate osmotic adaptations within seeds, including the accumulation of stress-responsive osmoregulators (Pamuru et al. 2021). It also disrupts the hormonal signalling, redox homeostasis and inhibits the cell elongation processes required for radicle emergence (Pamuru et al. 2021). Therefore, it is essential to improve seedling growth and plant development under drought stress to ensure crop productivity even under water-limited conditions.

Autophagy, one among several physiological processes modulates drought stress and ensures the survivability of the plants (Tang and Bassham 2021). This self-degradation involves the sequestration and degradation of non-functional cellular components, such as damaged organelles and proteins, within specialized vesicles called autophagosomes (Su et al. 2020; Wang et al. 2021). The controlled breakdown and recycling of these components provide energy and essential building blocks for cell survival and growth (Yang et al. 2019). Drought conditions were reported to enhance autophagy process by improving the expression of autophagy related genes (*ATGs*) and autophagosome formation. Additionally, *atg-*mutants showed hypersensitivity to drought (Liu et al. 2009). Seed autophagy is a finely orchestrated process that ensures the efficient mobilization of stored reserves, providing energy and nutrients essential for germination and vigorous seedling development (Iglesias-Fernández and Vicente-Carbajosa 2022; Wang et al. 2022).

Methylglyoxal (MGO) is a highly reactive toxic compound formed as a byproduct of glycolysis and other metabolic pathways, both enzymatically and non-enzymatically in plants (Allaman et al. 2015). It produced spontaneously via removal of phosphoryl group by β-elimination from 1, 2-enediolate of triose sugars dihydroxyacetone phosphate (DHAP) and glyceraldehyde-3-phosphate (G3P) (Hoque et al. 2012; Hossain et al. 2015). This dicarbonyl compound causes cellular damage by interacting with proteins, lipids, and deoxyribonucleic acid (DNA). Its toxicity is due to its high reactive nature resulting in the generation of reactive oxygen species (ROS) and formation of advanced glycation end products (AGEs), which inhibits germination and growth of plants (Hoque et al. 2012). Despite being toxic, recent research has highlighted the hormetic response of MGO as it functions as signalling molecule at lower intracellular levels by modulating various physiological processes in plants (Li 2016). Park et al. (2020) reported that MGO is a potent inducer of apoptosis whereas autophagy imparts protective role against MGO-induced cell death. Similarly, Lee et al. (2020) and Kim et al. (2020) highlighted that MGO regulates both autophagy and apoptosis within animal cellular systems. However, the precise interplay and connection between these intricately intertwined pathways remain to be comprehensively elucidated, particularly within the realm of plant systems.

Melatonin is well known for its role in regulating sleep and circadian rhythms in animals and has emerged as a significant player in plant biology influencing various physiological processes during stress conditions (Pan et al. 2023). Several reports have demonstrated that it combats abiotic stresses by regulating redox homeostasis and improving photosynthetic attributes. Li et al. (2018) reported that melatonin enhances heat tolerance of maize by detoxification of MGO, which helps in maintaining osmotic balance. Melatonin in combination with salicylic acid and gibberellic acid has shown to scavenge excess MGO by improving glyoxalase (GLY) system in wheat and tomato thus mitigating the effects of salinity stress (Siddiqui et al. 2020; Talaat and Todorova 2022). Melatonin is also known to have a cleaving effect on crosslinks in AGEs thus contributing to its degradation (Takabe et al 2016) and also prevents the AGE induced apoptosis *via* stimulation of autophagy flux (Jin et al. 2018).

Cotton (*Gossypium hirsutum* L.) is a globally important cash crop that yields fibre and oil, and plays an important role in India’s economic development. The significance of cotton extends beyond economics, encompassing cultural, social, and industrial domains (Zahid et al. 2021). However, the germination and yield potential of this ‘white gold’ is adversely affected by adverse environmental conditions, especially drought which is reported to alleviate germination rate up to 27.75% (Bai et al. 2020) and can decrease the yield up to 67% (Zafar et al. 2023). Drought disintegrates cellular harmony by accelerating the production of several toxic compounds such as methylglyoxal. Despite reports highlighting its potential to inhibit germination (Hoque et al. 2012), there have been studies stating that MGO enhances seed germination (Majláth et al. 2021). Therefore, the influence of drought-induced MGO production on the germination process is still obscure. Although, melatonin is already reported to improve seed germination under drought conditions (Bai et al. 2020), its regulatory effect on intracellular MGO homeostasis is yet to be explored. Additionally, drought-induced autophagy helps sustain the survivability of the plants (Liu et al. 2009), and therefore whether drought-induced MGO production and its homeostasis by melatonin can influence autophagy is still a question. Hence, the present research is directed towards shedding light on the involvement of melatonin in the modulation of intracellular MGO and its role in the induction of autophagy in both drought-tolerant and drought-sensitive cotton varieties under PEG induced drought stress.

## 2. Materials and methods

### 2.1. Seed materials and priming

Seeds of two varieties, *viz.,* L-799 (drought-sensitive) and Suraj (drought-tolerant) were obtained from Regional Agricultural Research Station, Guntur, Acharya N. G. Ranga Agricultural University (ANGRAU), Andhra Pradesh, India, and Central Institute for Cotton Research (CICR), Maharashtra, India, respectively. Seeds were surface sterilized with 3% (w/v) Bavistin (fungicide) for 5 min and 70% (v/v) ethanol for 3 min, followed by washing with autoclaved double distilled water. The seeds of two varieties were primed with different concentrations (5 µM, 10 µM, 25 µM, 50 µM, and 100 µM) of melatonin (Sigma-Aldrich, U.S.A.) by imbibing them for 24 h in the dark. The seeds imbibed in deionized water served as control. The seeds were then dried for 1 h at room temperature (RT) in the dark for back-drying and were evenly placed on moist sterile germination paper in germination boxes for 2 days in the dark at RT. Based on the initial experiments using different concentrations (10%, 15%, 20% and 25% w/v) of polyethylene glycol (PEG-6000), 18% PEG treatment for 4 days was selected for the study as it was optimal for providing stress, whereas 20% PEG resulting in the death of seedlings within 3 days while 15% PEG induced delayed stress responses. The two-day-old imbibed seeds were imposed drought stress by providing 18% (w/v) PEG and were allowed to grow in a culture room at 25 ± 2 °C, 65 ± 2% relative humidity under 16:8-h light and dark photoperiod for another 4 days. The 6-day old seedlings were used to carry out all the experiments with three independent set of plants, with triplicates per treatment in each replication of plants.

### 2.2. Measurement of RWC, ROS, lipid peroxidation and electrolyte leakage

Relative water content (RWC) was estimated using the method described by Weatherley (1950) method and expressed in percentage (%).

Superoxide anion radical (O_a_^−^) and hydrogen peroxide (H_2_O_2_) were histochemically detected using nitrogen blue tetrazolium (NBT, 1 mg. ml^−1^) and 3,3’-diaminobenzidine stains (DAB, 0.5 mg.ml^−1^) as described by Ramel et al. (2009). The H_2_O_2_ content was determined according to the method described by Okuda et al. (1991). Approximately 100 mg of seedlings were homogenized in 0.1% trichloroacetic acid (TCA) solution and centrifuged at 12,000 *g* for 15 min. The supernatant was collected and mixed with 10 mM potassium phosphate buffer (pH-7.0) and 1 M potassium iodide in 1:1:2 ratio and absorbance was taken immediately at 390 nm. The extinction coefficient of H_2_O_2_ produced is 0.28 µmol.cm^−1^ and expressed in µmol.gm^−1^.

Lipid peroxidation was determined by measuring malonaldehyde (MDA) content (Heath and Packer 1968). Seedlings were homogenized by adding 0.1 % (^w^_/v_) TCA followed by centrifugation at 12000 *g* at 4 °C for 10 min. The supernatants were collected and mixed with 0.5% (^w^_/v_) thiobarbituric acid (ΤΒΑ) diluted in 20 % (^w^_/v_ ) TCA in 1:3 ratio and incubated at 95 °C for 30 min in the water bath. The reaction was terminated by incubating on ice. The absorbance was measured at 532 and 600 nm. Optical density (OD_600_) value was subtracted from OD_532_, and MDA concentration was calculated using the Lambert-Beer law with an extinction coefficient εΜ= 155 mM^−1^ cm^−1^.

Electrolyte leakage (EL) was estimated following the protocol of Dionisio-Sese and Tobita (1998) with modification. The seedlings were immersed in deionized water in a test tube and heated for 2 h at 25 °C, and their conductivity was measured (EC_a_). Then, the tubes with the seedlings were heated at 100 °C for 15 min, and after cooling them to RT, the conductivity was measured (EC_b_).

### 2.3. Protein extraction and enzymatic assays

Crude protein was isolated to determine enzyme activities. Protein was extracted from fresh seedlings (200 mg) with extraction buffer containing 100 mM sodium phosphate buffer (pH-7.0), 50% glycerol, 16 mM magnesium sulfate (MgSO_4_), 0.2 mM phenylmethylsulfonyl fluoride (PMSF) and 0.2% (^w^_/v_) polyvinylpolypyrrolidone (PVPP). The extracted protein was quantified employing the Lowry method (Lowry et al. 1951) and used in enzyme assays.

The GLY I (EC 4.4.1.5) and GLY II (EC 3.1.2.6) enzyme activities were determined by measuring the generation and utilization of S-lactoylglutathione (SLG) (Singla-Pareek et al. 2003; Sahoo et al. 2021). To determine GLY I activity, the reaction mixture containing 100 mM sodium phosphate buffer, pH 7.5, 3.5 mM MGO, 1.7 mM glutathione (GSH) and 16 mM MgSO_4_ was incubated in the dark for 7 min which forms the intermediate substrate hemithioacetal. After incubation, 25 μg of crude protein extract was added into the 1 ml of reaction mixture and absorbance was taken at 240 nm within a timescale that showed linear increase in the absorbance. The molar extinction coefficient of generated SLG is 3370 M^−1^ cm^−1^ and expressed in µmol min^−1^ mg^−1^ protein. To determine GLY II activity, 25 μg of crude protein was added into buffer containing 50 mM 3-(N-morpholino) propanesulfonic acid (MOPS, pH-7.2), 0.6 mM SLG and the decrease in absorbance was measured at 240 nm. The molar extinction coefficient of generated D-Lactate is 3.1 mM^−1^ cm^−1^ and expressed in µmol min^−1^ mg^−1^ protein.

GLY III (EC 4.2.1.130) enzyme activity was measured by colorimetric method proposed by Ghosh et al. (2022) with slight modifications. Total protein of 20 μg was added to 10 mM sodium phosphate buffer (pH-7.4), 1 μM of MGO and incubated for 15 min at 45 °C. Then, 2,4-dinitrophenylhydrazine (DNPH) solution was added and incubated for another 15 min at RT, and the absorbance was measured at 550 nm with a UV-Vis (Ultraviolet-Visible) spectrophotometer (Shimadzu, Japan). The buffer with MGO and DNPH solution devoid of protein was used as a control. Specific activity was calculated by subtracting the final MGO content from the initial one and expressed in nmol min^−1^ mg^−1^ of protein.

### 2.4. Determination of glutathione pool

Total glutathione was measured according to the method described by Sahoo et al. (2017). The seedlings were macerated, homogenised in potassium phosphate buffer (0.5 M, pH 7.5) and centrifuged at 10000 *g* for 15 min at 4 °C. The resulting supernatant (1 ml) was added to 100 µl 2-nitrobenzoic acid (DTNB, 10 mM), 200 µl bovine serum albumin (BSA, 10 mM), 100 µl nicotinamide adenine dinucleotide (NADH, 0.5 mM) and incubated at 37 °C for 15 min. The absorbance was taken at 412 nm and expressed as μmol g^−1^ fresh weight (FW).

For determination of oxidized glutathione (GSSG), the GSH was masked by adding 2-vinylpyridine to the supernatant and incubated for 1 h at 25 °C. The extract (100 µl) was combined with 600 µl reaction buffer (100 mM potassium phosphate buffer and 5 mM EDTA, pH 7.5), 100 µl of glutathione reductase (GR, 20 U.ml^−1^), and 100 µl of DNTB (10 mM). The reaction was initiated by adding 100 µl of nicotinamide adenine dinucleotide phosphate NADPH (2.5 mM), and the rate of absorption was measured at 412 nm. The results are expressed as nmol *g*^−1^ FW. The reduced glutathione (GSH) content was determined by subtracting GSSG from total glutathione.

### 2.5. Estimation of MGO content

Methylglyoxal was estimated as described previously by Ghosh et al. (2014). Approximately 300 mg of seedlings were macerated with liquid nitrogen, homogenized in 2.5 ml perchloric acid (0.5 M) and centrifuged after 5 min at 12000 *g* for 10 min at 4 °C. The resulting supernatant, mixed with charcoal (10 mg ml^−1^) for decolorization, was neutralized with 1 M sodium hydrogen phosphate (Na_2_HPO_4_). For quantification, a reaction mixture was prepared using 250 μl of 7.2 mM 1,2-diaminobenzene, 100 μl of 5 M perchloric acid, 10 μl of 100 mM sodium azide and 650 μl of neutralized supernatant. After incubating at RT for 3 h, the absorbance was taken at 336 nm. A standard curve was plotted using various concentrations of MGO (0, 5, 10, 25, 50 and 100 μM), and the content was expressed as μmol g^−1^ FW.

### 2.6. Quantification of glucose content

Total sugars were extracted from seedlings following the method of Giannoccaro et al. (2006) and detected using reverse phase high-performance liquid chromatography (HPLC) (LC-20AD Shimadzu, Japan). Approximately. 100 mg of seedlings were ground in 1 ml of milliQ water in a 1:10 (w/v) ratio for 15 min in a rotospin. After centrifugation at 12000 *g* for 10 min at RT, 500 µl of the clear supernatant was mixed with 1.5 ml of 95% acetonitrile for 30 min in the rotospin. The resulting supernatant was collected and evaporated in a dry bath at 95 °C. The residue was re-dissolved in 1 ml of milliQ water and filtered through a 0.22-µm filter paper using syringe filters. Different sugars were separated isocratically through reverse phase HPLC using an amino (NH_2_) column with acetonitrile and water (70:30) as the mobile phase. The flow rate was set to 1 ml.min^−1^ and absorbance was detected at 190 nm (UV) using a photodiode array (PDA) detector. The glucose peak was identified by spiking with standards and concentrations were calculated using an external standard calibration method (Sreeharsha et al. 2018).

### 2.7. Determination of pyruvate content

The pyruvate content was quantified by homogenising 500 mg of seedlings in 1 ml water, which was allowed to stand for 10 min at RT, and centrifuge at 10000 g for 5 min. To the resulting supernatant, 1 ml of 0.25 g l^−1^ DNPH in 1 M hydrochloric acid (HCl) was added and placed in a water bath at 37 °C for 10 min and 1 ml of 1.5 M sodium hydroxide (NaOH) was added. Then, 1 ml of 1.5 M NaOH was added, and absorbance was measured at 515 nm. Standards were prepared by adding 25–200 µl of 1 mM sodium pyruvate and reducing the amount of water in the assay accordingly (Anthon and Barrett 2003).

### 2.8. Quantification of endogenous melatonin content

Seedling samples were weighed, sliced into small (3–5 mm) pieces and immersed into chloroform vials, followed by overnight shaking at 4 °C. The seedling pieces were removed, and the solvent in the vial was evaporated under nitrogen (N_2_) gas at 4 °C. The resulting residue was dissolved in acetonitrile, filtered through a 0.2 µm polyvinylidene difluoride (PVDF) membrane filter, and used for HPLC analysis. Shimadzu HPLC system (Kyoto, Japan) equipped with a C18 column (Phenomenex KINETEX, 250 mm × 4.6 mm) was used to determine the melatonin content. The mobile phase used was a 50:50 mixture of water and acetonitrile with a flow rate of 1 ml min^−1^ and detection was performed at 280 nm using UV. The melatonin levels were quantified using pure melatonin (Sigma-Aldrich, U.S.A.) as a standard, following the method Arnao and Hernández-Ruiz (2014) described.

### 2.9. Determination of anti-glycation activity

The assay for estimating the inhibition of AGE formation was performed following the method outlined by Peng et al. (2008). Methylglyoxal and bovine serum albumin (BSA) were mixed in 100 mM potassium phosphate (pH-7.4) to make a final concentration of 5 mM and 1 mg ml^−1^, respectively. After the addition of sodium azide (0.2 g l^−1^) the mixture was incubated at 37 °C for 5 days with or without crude protein extracts (20 μg). Phosphate buffer was used as a blank, and aminoguanidine (5 mM), a known AGE inhibitor and MGO scavenger, was used as positive control. The fluorescence was measured at 340 nm excitation and 420 nm emission (Ghosh et al. 2022).

### 2.10. Measurement of the loss of cell membrane integrity

Evans blue, an anionic molecule, selectively permeates only ruptured plasma membranes. To quantitatively assess dye absorption per unit dry weight of plant samples, the seedlings were immersed in a 0.5% (w/v) Evans blue solution and incubated for 15 min at RT under vacuum. Subsequently, the samples were transferred to a destaining solution and incubated at 60 °C for 30 min to 1 h. The absorption of supernatant was measured at 600 nm using the destaining solution as a blank, and the pellet was dried at 60 °C overnight, and the weight was determined. The A600 values were normalized to dry weight (Minina et al. 2013).

### 2.11. Quantification of ATP

The adenosine triphosphate (ATP) content in the seedlings was measured using the protocol described by Zhang et al. (2020). Approximately 10 mg of leaf tissue was powdered in liquid N_2_, followed by homogenization in 100 µl of 0.1 M HCl. The homogenate was then combined with 820 µl of buffer solution (1 M citric acid and 1 M di-sodium hydrogen phosphate) along with 80 µl of chloroacetaldehyde. The reaction mixture was incubated for 10 min at 80 °C and subsequently centrifuged (12000 g, 10 min). The resulting supernatant was filtered through 0.25 µm filter paper. The filtrate was then subjected to HPLC analysis, and Adenosine 5′-triphosphate disodium salt hydrate (Sigma, U.S.A) was used as the standard.

### 2.12. Determination of autophagosomes and plant cell viability

Monodansylcadaverine (MDC) staining was used to detect autophagy following the protocol of Contento et al. (2005). Seedlings were incubated in 0.05 mM MDC (Sigma Aldrich, U.S.A.) in phosphate-buffered saline (PBS) for 10 min, followed by three washes with PBS at RT. Seedlings were imaged using a Leica SP8 laser scanning confocal microscope (NLO 710, Carl Zeiss, Germany) using a 4′,6-diamidino-2-phenylin-dole (DAPI)-specific filter. The excitation and emission wavelengths for MDC were 345 and 455 nm, respectively. Fluorescein diacetate (FDA) staining was used to detect cell viability; the seedlings were stained with 5 µg ml-1 (FDA; Sigma) for 5 min and were washed three times using sterile water (Guan et al. 2019). Fluorescence signals were visualized on confocal microscope with an excitation and emission wavelengths of 488 nm and 525 nm. The quantitative measurement of fluorescence intensity was carried out using ImageJ.

### 2.13. Total protein extraction and immunoblot analysis

The seedlings were macerated in liquid N_2_ and homogenized in a buffer containing 50 mM Tris– HCl (pH-8.0), 150 mM sodium chloride (NaCl), 1 mM phenyl-methanesulfonyl fluoride, and 10 mM iodoacetamide (Chung et al. 2009) and protease inhibitor cocktail. For immunoblot analysis of ATG8, 15% of SDS-PAGE gel was prepared with 6 M urea (Wang et al. 2019) and for MAPK6 (Mitogen Activated Protein Kinase 6) and KIN10 (Snf1-Related Kinase homolog 10), 10% of SDS-PAGE gels were prepared. Following electrophoresis, the protein-containing SDS-PAGE gels were transferred to a nitrocellulose membrane. ATG8 and ATG8-PE levels were determined using an anti-ATG8 antibody (AS14 2769, Agrisera, Sweden) in 1: 500 dilution, anti-MPK6 (AS12 2633, Agrisera, Sewden) in 1: 250 and anti-SnRK1alpha1 (AS214581, Agrisera, Sweden) in 1:500. Histone H3 was used as a loading control. Thus, for determining the His-H3 level, anti-histone-H3 (AS10710, Agrisera, Sweden) was used as primary antibody in 1:2000 dilution. An anti-rabbit antibody conjugated to HRP was used as the secondary antibody.

### 2.14. Quantification of ABA

The ABA content was quantified using Liquid Chromatography-Mass Spectrometry/Mass Spectrometry (LC-MS/MS). Samples were prepared according to the method outlined by Pan et al. (2010). The analysis was conducted on an Agilent Quadrupole Time of Flight (Q-TOF) LC/MS 6520 series system (Agilent Technologies, U.S.A.) equipped with a ZORBAX RX-C18 column (4.6 × 150 mm, 5 μm, Agilent Technologies, U.S.A.) at 24 °C, as described in the report of Shreya *et al*. (2022). The mass detection range was set between 100 and 2000 m z^−1^.

### 2.15. RNA Isolation, cDNA Synthesis, and quantitative real-time PCR

Total ribonucleic acid (RNA) was isolated using the cetyltrimethylammonium bromide (CTAB)-ammonium acetate method (Zhao et al. 2012). RNA quality was assessed through gel electrophoresis and nanodrop 2000 UV-Vis spectrophotometer (Thermo Scientific, U.S.A.). Primers targeted for the genes were synthesized using GenScript1 (**Supplementary table 1**). Complementary deoxyribonucleic acid (cDNA) was synthesized from the total ribonucleic acid (RNA) using primescript 1^st^ strand synthesis Kit (Takara Bio Inc., Japan) and Real-time polymerase chain reaction (PCR) was performed on mastercycler realplex (Eppendorf, Germany). Actin4 was used as an internal control, and the relative fold-change RNA expression was estimated using the 2^−ΔΔCT^ method (Livak and Schmittgen 2001).

### 2.16. Statistical analysis

The data represent the mean values of 3 independent treatments, with 3 replicates per treatment in each experiment. These data were subjected to one-way analysis of variance (ANOVA) using SigmaPlot (12.0 version) software. The error bars depicted in the graph represent the standard error (± SE) of the mean values. To evaluate the significance of differences between treatments, Duncan’s multiple range test (DMRT) was performed at a significance level (*p* ≤ 0.05). All the graphs were made using PRISM8 and Figure 9 in BioRender software.

## 3. Results

### 3.1. Effect of melatonin on seedling phenotype and RWC under drought stress

The effects of melatonin priming on the growth of the seedlings varied in drought distinguished varieties depending on the concentrations used. The seeds of L-799 primed with 25 µM melatonin showcased improved growth under PEG-induced drought stress, surpassing other concentrations. This concentration also resulted in the highest radicle length among all the primed control seedlings (**Figure 1A and Supplementary figure 1A**). Contrarily, in Suraj, seeds primed with melatonin at all applied concentrations did not cause distinct phenotypic changes in the seedlings compared to unprimed seedlings under stress (**Figure 1A and Supplementary figure 1B**). Therefore, 25 µM concentration of melatonin was chosen for comparative studies in both the varieties.

**Figure 1:**
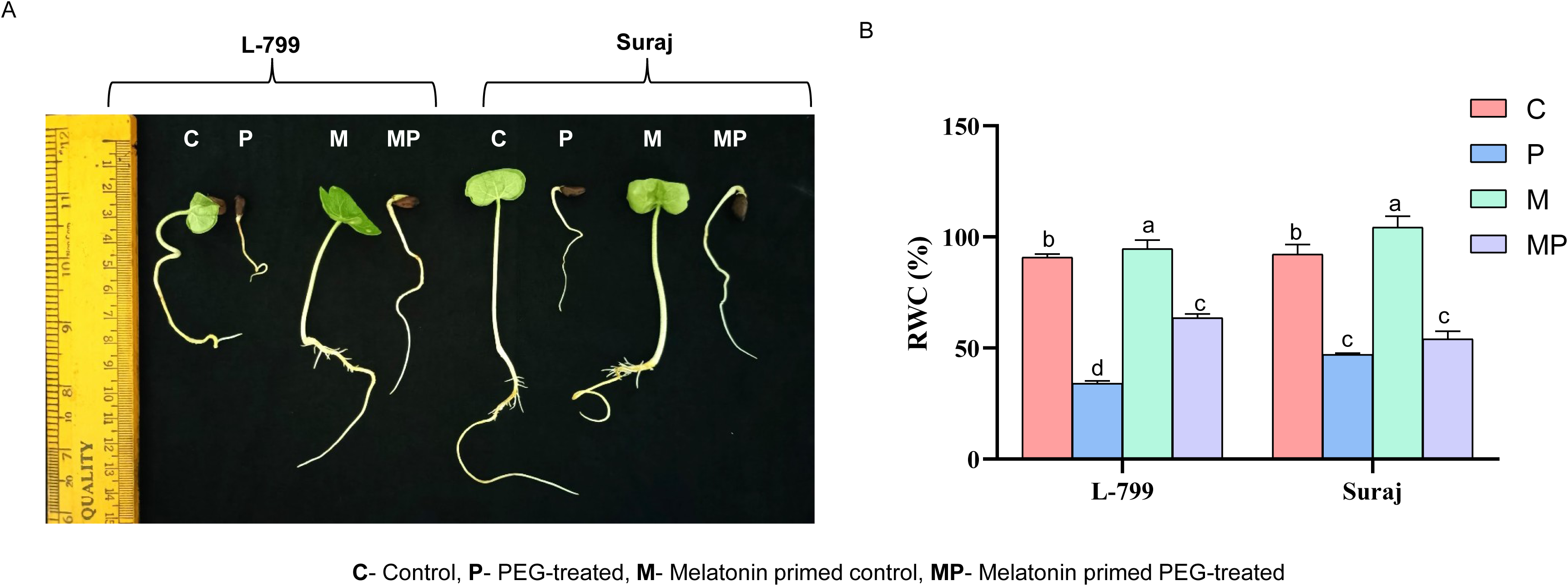
Effect of melatonin priming on seedling growth and relative water content (RWC) under optimal and PEG-induced stress conditions for 4 days in drought-sensitive (L-799) and drought-tolerant (Suraj) varieties of cotton (*Gossypium hirsutum* L.). (**A**) Phenotype of the and (**B**) RWC. The data represent mean values ± SE of 3 independent experiments, with 3 replicates per treatment in each experiment. Different alphabets within the group represent significant differences among the treatments according to Duncan’s multiple range test (DMRT) at *p*-value ≤ 0.05.

The relative water content (RWC), an indicator of cellular turgor pressure, reflects the plant’s water status and capacity to maintain turgor. Melatonin-primed L-799 exhibited significantly higher RWC (86.82%) compared to unprimed seedlings during drought stress. Conversely, Suraj showcased no significant difference in RWC between primed and unprimed seedlings under stress (**Figure 1B**).

### 3.2. Influence of exogenous melatonin priming on endogenous melatonin content

Exogenous melatonin priming is reported to increase the endogenous melatonin content in cotton under drought stress (Supriya et al. 2022). The seedlings of unprimed stressed L-799 showed a 1.11-times decreased endogenous melatonin compared to the controls, whereas in primed seedlings it increased by 4.87-times than unprimed under stress. Intriguingly, unprimed Suraj seedlings under stress exhibited 2.31-times higher melatonin content than the controls. On the other hand, primed Suraj showed a decrease in melatonin by 1.11-times compared to unprimed seedlings under stress (**Figure 2A**).

**Figure 2:**
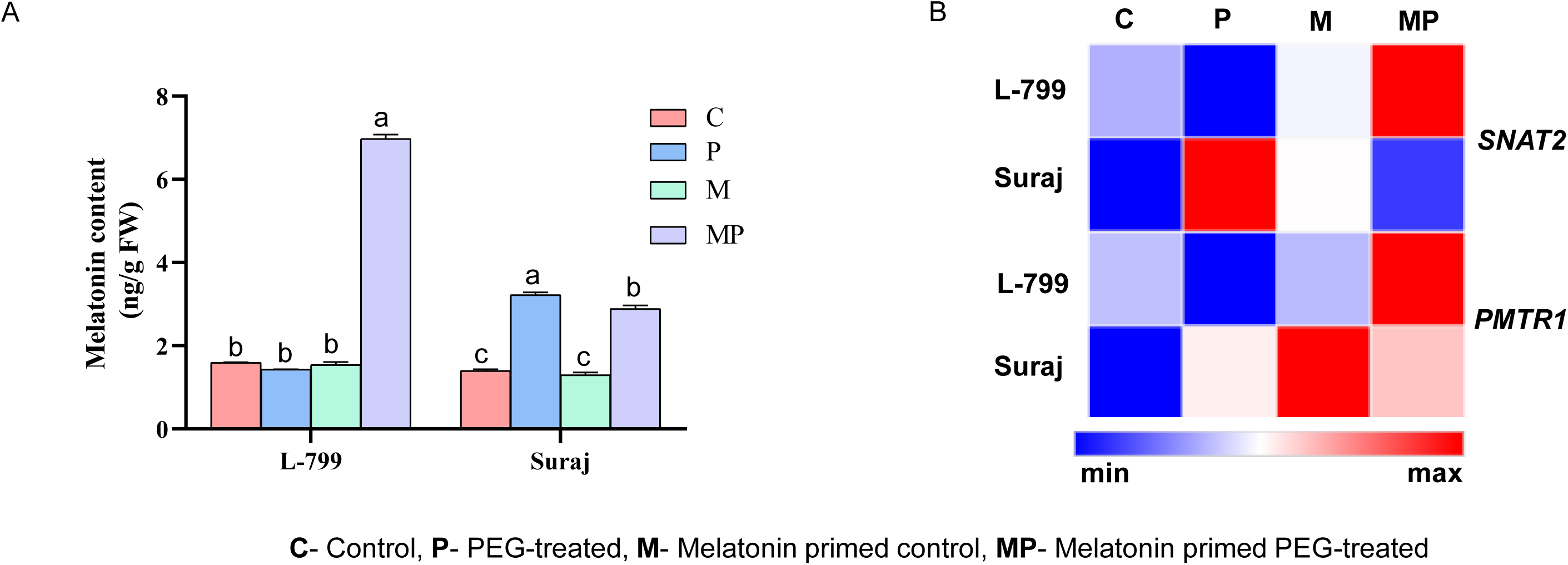
Effect of melatonin priming on endogenous melatonin content and expression analysis of genes of melatonin biosynthesis and signalling under optimal and PEG-induced stress conditions in drought-sensitive (L-799) and drought-tolerant (Suraj) varieties of cotton (*Gossypium hirsutum* L.). (**A**) Melatonin content, and (**B**) Heatmap of relative fold-change expression of melatonin biosynthesis gene (*SNAT2*) and melatonin signalling receptor gene (*PMTR1*). The data represent mean values ± SE of 3 independent experiments, with 3 replicates per treatment in each experiment. Different alphabets represent significant differences among the treatments according to Duncan’s multiple range test (DMRT) at *p*-value ≤ 0.05.

The expression of the melatonin biosynthesis gene, serotonin N-acetyl transferase 2 (*SNAT2*), was downregulated by 17.54-fold in unprimed stressed seedlings compared to controls. However, priming caused significant upregulation (98.59-fold) compared to unprimed in L-799 under stress. Contrastingly, in Suraj, although unprimed stressed seedlings exhibited significantly enhanced expression of *SNAT2* (10.94-fold) compared to controls, the primed seedlings showed 7.10-fold reduced expression compared to unprimed under stress conditions. Furthermore, the expression levels of the *phytomelatonin receptor 1* (*PMTR1*) were notably increased in both L-799 (1.91-fold) and Suraj (1.07-fold) in melatonin-primed seedlings compared to respective unprimed seedlings under drought stress (**Figure 2B, Supplementary figure 2A and B**).

### 3.3. Role of melatonin on intracellular ROS accumulation, lipid peroxidation and electrolyte leakage under drought stress

During stress conditions, unprimed plants exhibited overaccumulation of O_2_^−^ and H_2_O_2_, as reflected by intense NBT (**Figure 3A**) and DAB staining (**Figure 3B**). However, priming effectively reduced ROS accumulation compared to unprimed plants in both varieties. The intracellular H_2_O_2_ decreased by 25.31% and 26.69% in primed seedlings, compared to respective unprimed under stress in L-799 and Suraj varieties, respectively. (**Figure 3C**).

**Figure 3:**
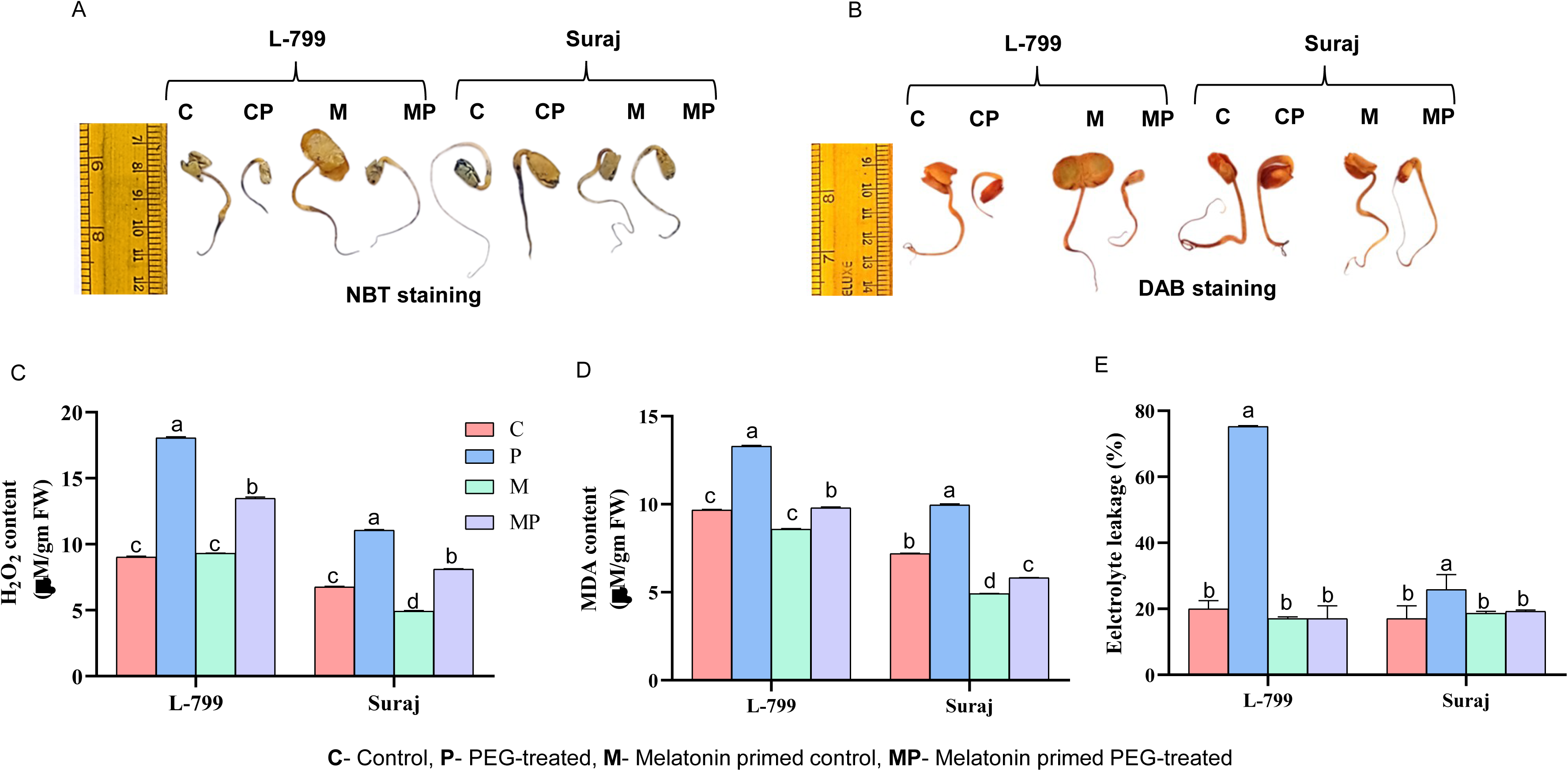
Effect of melatonin priming on ROS, MDA content and EL under optimal and PEG-induced stress conditions for 4 days in drought-sensitive (L-799) and drought-tolerant (Suraj) varieties of cotton (*Gossypium hirsutum* L.). (**A**) Histochemical detection of (O_2_^−^) through NBT staining, (**B**) Histochemical detection of (H_2_O_2_) through DAB staining, (**C**) Quantification of H_2_O_2_ content, (**D**) MDA content and (**E**) EL. The data represent mean values ± SE of 3 independent experiments, with 3 replicates per treatment in each experiment. Different alphabets represent significant differences among the treatments according to Duncan’s multiple range test (DMRT) at *p*-value ≤ 0.05.

Lipid peroxidation disrupts membrane structure, thereby exhibiting deleterious effects on the function of the cells. Both L-799 and Suraj showed 37.43% and 38.38% elevated MDA content in unprimed stressed seedlings compared to controls, but priming significantly lessened the content by 26.33% and 41.50% compared to unprimed under stress conditions, respectively (**Figure 3D**). Electrolyte leakage (EL), a consequence of lipid peroxidation, was significantly increased in both unprimed L-799 (3.76 times) and Suraj (1.51 times) under stress compared to controls. Priming reversed the scenario by decreasing EL by 4.41 and 1.34 times in both varieties compared to respective unprimed seedlings under stress (**Figure 3E**).

### 3.4. Effect of melatonin on intracellular MGO levels

The MGO content was significantly increased (9.39 times) in L-799 unprimed stressed seedlings compared to controls, whereas priming decreased the content significantly by 7.23-times compared to unprimed seedlings under stress. Interestingly, unprimed stressed seedlings of Suraj exhibited a notable decrease in MGO content (2.6-times) compared to controls but displayed no significant change in primed and unprimed stressed seedlings (**Figure 4A**).

**Figure 4:**
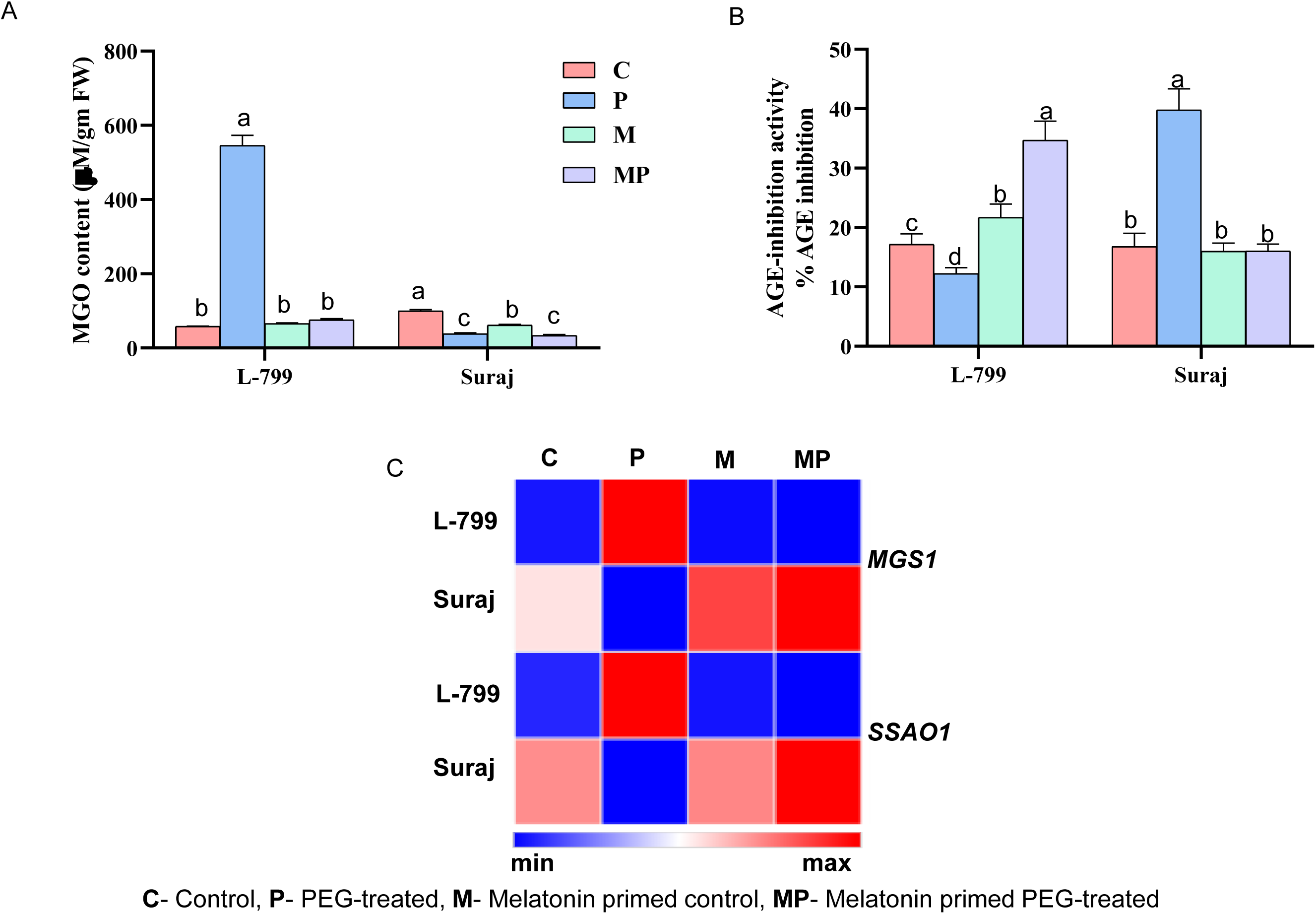
Effect of melatonin priming on endogenous MGO content, expression of MGO biosynthesis genes and AGE formation under optimal and PEG-induced stress conditions in drought-sensitive (L-799) and drought-tolerant (Suraj) varieties of cotton (*Gossypium hirsutum* L.). (**A**) MGO content, (**B**) Anti-glycation activity, and (**C**) Heatmap of relative fold change expression of *MGS1* and *SSAO1*. The data represent mean values ± SE of 3 independent experiments, with 3 replicates per treatment in each experiment. Different alphabets within the group represent significant differences among the treatments according to Duncan’s multiple range test (DMRT) at *p*-value ≤ 0.05.

MGO is known to form AGEs in the cell. The AGE inhibition (%) activity decreased by 1.4 times in unprimed stressed seedlings compared to controls, while priming elevated the activity by 2.83 times compared to unprimed in L-799 under stress. Interestingly, unprimed Suraj exhibited 2.36 times enhanced activity compared to controls, but priming decreased the activity by 2.48 times compared to unprimed seedlings under stress (**Figure 4B**).

Methylglyoxal synthase (MGS) and semicarbazide-sensitive amine oxidase (SSAO) genes are involved in MGO biosynthesis. The transcript levels of both *MGS1* and *SSAO1* increased significantly by 33.89 and 18.20-fold in unprimed stressed seedlings, respectively, compared to controls in L-799. However, priming significantly alleviated the expressions by 178.36 and 67.40-fold compared to the unprimed seedlings under stress conditions. However, unprimed stressed seedlings of Suraj displayed downregulated expressions of both *MGS1* (2.50-fold) and *SSAO1* (2.85-fold) compared to controls. However, Suraj primed seedlings showed 5.12- and 4.08-fold increased expression of *MGS1* and *SSAO1* compared to unprimed seedlings under stress, respectively (**Figure 4C Supplementary figure 2C and D**).

### 3.5. Effect of melatonin priming on glyoxalase and non-glyoxalase systems

Glyoxalase I, II and III are enzymes involved in the MGO detoxification pathway. The activities of these enzymes decreased by 1.12, 4.64 and 5.03 times in unprimed stressed seedlings compared to controls in L-799. Priming caused an increase in the activities by 1.70, 6.64 and 22.33 times compared to unprimed seedlings, respectively during stress. In Suraj, the activities increased by 2.58,

1.77 and 2.67-times in unprimed stressed compared to control seedlings, but it decreased significantly upon priming by 1.30, 1.47 and 2.59 times compared to unprimed seedlings under stress **(Figure 5A-C)**. Additionally, the transcript levels of *GLY I-1*, *II-2* and *III-DJ-1 homolog* were downregulated (20.40, 20.30, and 12.50-fold, respectively) in unprimed L-799, whereas they were upregulated (11.43, 9.19, and 10.09-fold, respectively) in unprimed Suraj, compared to respective controls under stress. Priming reversed the scenario, as primed L-799 seedlings showed upregulated expression (63.46, 63.12, and 84.37-fold), while it decreased (30.89, 3.34, and 2.29-fold) in primed Suraj, compared to respective unprimed seedlings under drought (**Figure 5D and Supplementary figure 3A-C**).

**Figure 5:**
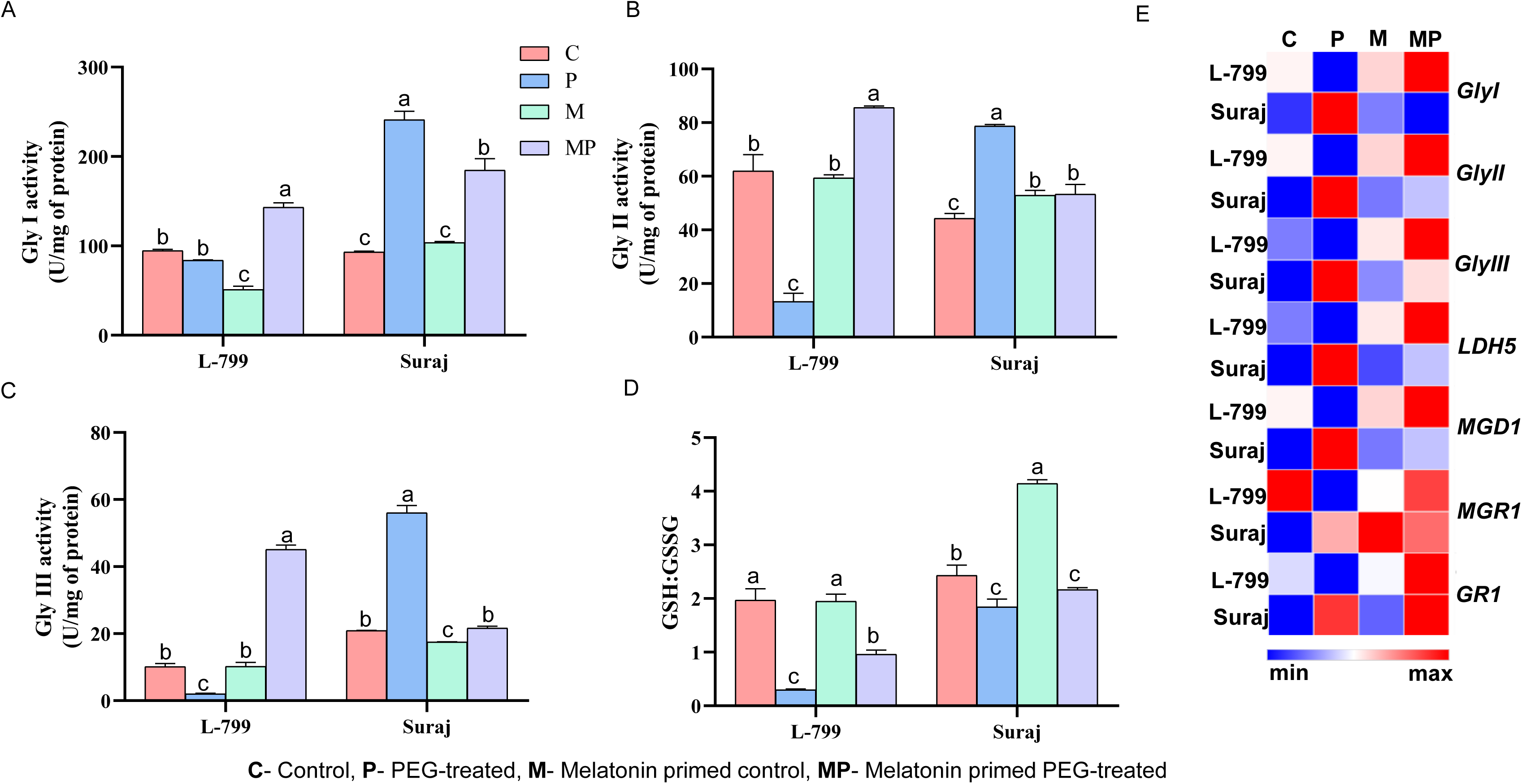
Effect of melatonin priming on glyoxalase, non-glyoxalase system and glutathione pool under optimal and PEG-induced stress conditions in drought-sensitive (L-799) and drought-tolerant (Suraj) varieties of cotton (*Gossypium hirsutum* L.). (**A**) Gly I activity, (**B**) Gly II activity, (**C**) Gly III activity, (**D**) GSH:GSSG, and (**E**) Relative fold-change expression of glyoxalase system genes *Gly I*, *Gly II* and *Gly III ,* non-glyoxalase system genes *LDH5*, *MGD1* and *MGR1* and glutathione reductase gene (*GR1*). The data represent mean values ± SE of 3 independent experiments, with 3 replicates per treatment in each experiment. Different alphabets represent significant differences among the treatments according to Duncan’s multiple range test (DMRT) at *p*-value ≤ 0.05.

Lactate dehydrogenase (LDH), methylglyoxal dehydrogenase (MGD), and methylglyoxal reductase (MGR) are genes of the non-glyoxalase system involved in MGO scavenging. The expression of these genes was downregulated by 12.50, 25 and 2-fold in unprimed stressed seedlings but were upregulated (84.37, 77.75 and 1.80-fold, respectively) in primed stressed seedlings compared to unprimed in L-799. In contrast, Suraj displayed an upregulated expression of 10.09, 9.19, and 1.63-fold in unprimed stress compared to controls, respectively. Although priming upregulated the *MGR1* expression (1.13-fold), the expression of *LDH5* (2.29-fold) and *MGD1* (3.34-fold) were downregulated significantly, compared to unprimed seedlings under stress (**Figure 5E and Supplementary figure 3 D-F**).

### 3.6. Effect of melatonin on glutathione level

Glutathione serves as a fundamental antioxidant, and the GSH/GSSG ratio plays a crucial role in maintaining the redox homeostasis of plants under stress. Melatonin priming elevated the GSH levels (3.03 times), GSH/GSSG ratio (3.31 times) and total glutathione (1.42 times) in L-799 seedlings compared to unprimed under stress. However, Suraj showed no notable changes in these attributes in primed and unprimed seedlings under stress. However, priming decreased GSSG content by 1.05 times and 1.10 times in L-799 and Suraj seedlings compared to unprimed under stress (**Figure 5D and Supplementary figure 4A-C**). Furthermore, priming elevated *GR1* expression by 241.76-fold in L-799 and 1.16-fold in Suraj, compared to respective unprimed seedlings under stress (**Figure 5E and Supplementary figure 4D**).

### 3.7. Effect of melatonin on glycolysis and related enzymes under stress

In primed L-799 seedlings, the expression of glycolysis-related genes showed distinct changes, with a 3.46-fold decrease in hexokinase 3 (*HK3*) and a 2.7-fold increase in triosephosphate isomerase (*TPI1*) expression compared to unprimed seedlings under stress. Unlike L-799, unprimed Suraj displayed a 1.52-fold decreased *HK3* and a substantial increase (12.11-fold) in the expression of *TPI1* compared to primed seedlings under stress. Furthermore, priming upregulated the expression of phosphoglycerate kinase 5 (*PGK5*) and pyruvate kinase 1 (*PK1*) (9.68 and 31.33-fold, respectively) in L-799, whereas downregulated them (2.98 and 1.21-fold, respectively) in Suraj compared to unprimed seedlings under stress conditions (**Figure 6A and Supplementary figure 5A-D**).

**Figure 6:**
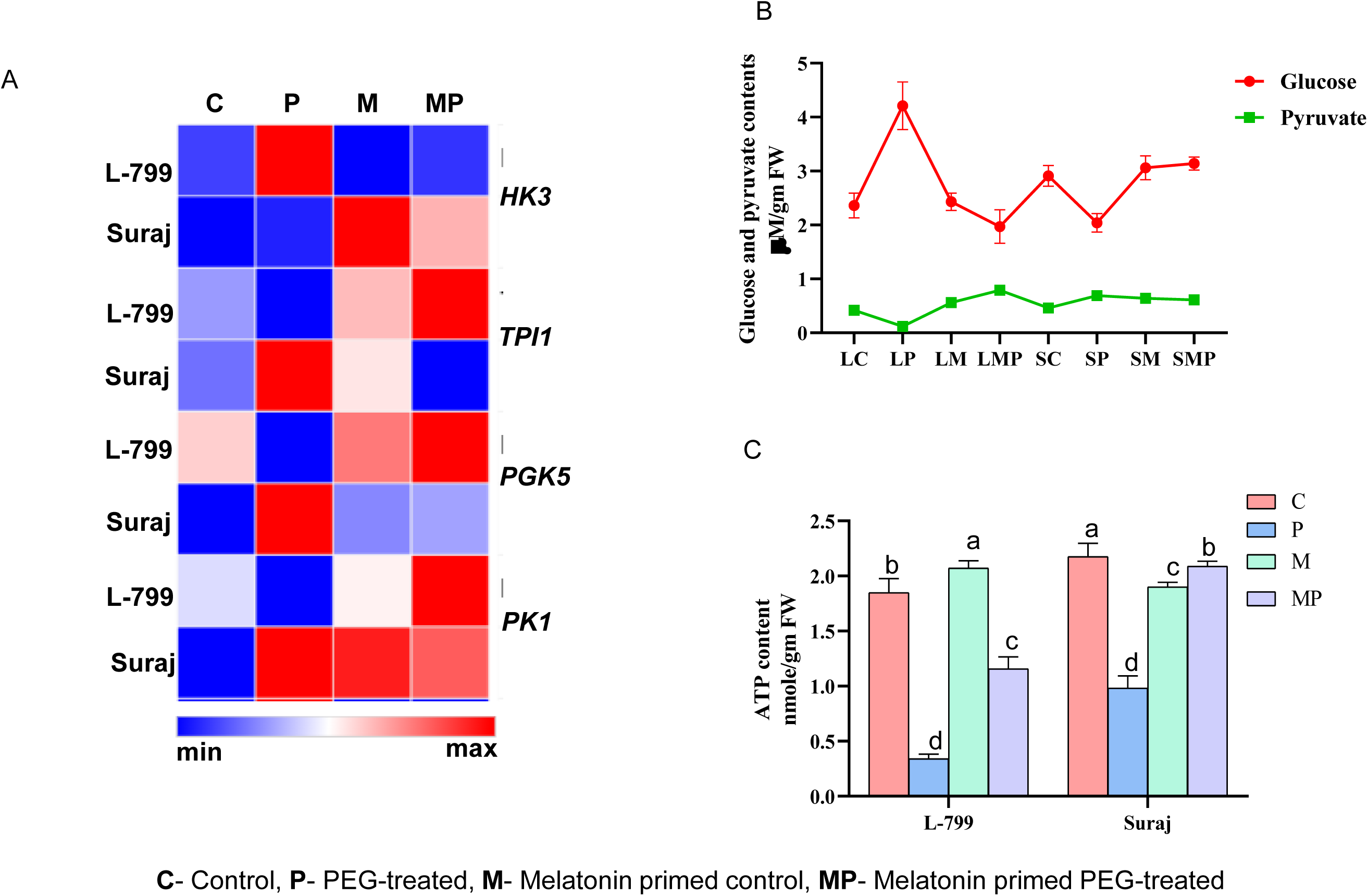
Effect of melatonin priming on glycolysis and expression of related genes under optimal and PEG-induced stress conditions for 4 days in drought-sensitive (L-799) and drought-tolerant (Suraj) varieties of cotton (*Gossypium hirsutum* L.). (**A**) Relative fold-change expression of glycolysis related genes *HK3, TPI1, PGK1* and *PK1,* (**B**) Endogenous glucose content and pyruvate content, and (**C**) ATP content. The data represent mean values ± SE of 3 independent experiments, with 3 replicates per treatment in each experiment. Different alphabets represent significant differences among the treatments according to Duncan’s multiple range test (DMRT) at *p*-value ≤ 0.05.

The intracellular glucose content was elevated (1.78 times) in unprimed stressed seedlings of L-799, but priming reduced it (2.13 times) compared to unprimed seedlings under stress. However, Suraj’s unprimed stressed seedlings displayed significantly decreased glucose content (1.42 times) compared to controls, whereas priming decreased it by 1.53 times compared to unprimed seedlings under stress. The intracellular pyruvate content decreased by 3.5 times in unprimed L-799 but was elevated (1.50 times) in unprimed Suraj under stress compared to respective control seedlings. Priming reversed the scenario as pyruvate levels increased 6.58times in L-799 but decreased by 1.13 times in Suraj compared to respective unprimed seedlings under stress (**Figure 6B, Supplementary figure 5E and F**).

### 3.8. Effect of melatonin on ATP level

Stress conditions are known to reduce the intracellular ATP content. In this study, stress conditions alleviated the intracellular ATP level in unprimed seedlings of both L-799 (5.44 times) and Suraj (2.21 times) compared to respective controls. However, priming improved the level significantly by 3.40 (in L-799) and 2.15 times (in Suraj) compared to respective unprimed seedlings under stress **(Figure 6C)**.

### 3.9. Regulation of autophagy under drought stress by melatonin

Melatonin is known to regulate autophagy to sustain the growth and survivability of plants under stress conditions and is reflected in the increased expression of *ATG* genes and autophagosome formation. Lesser MDC-stained bodies (autophagosomes) with notably downregulated expressions of *ATG8c* (3.70-fold), *ATG3* (3.70-fold), *ATG5* (4-fold) and *ATG18a* (3.70-fold) in unprimed stressed compared to controls in L-799 indicates reduced autophagosome formation. Priming reversed the situation, as evident from increased autophagosome structure (MDC-stained structure) with upregulated expression of *ATG8c* (12.55-fold), *ATG3* (12.81-fold), *ATG5* (9.16-fold) and *ATG18a* (12.37-fold) in primed seedlings compared to unprimed under stress. Unlike L-799, the transcript levels of the autophagy-related genes were significantly higher in unprimed stressed compared to primed seedlings of Suraj. However, no significant difference was observed in MDC-stained bodies in primed and unprimed stressed Suraj seedlings **(Figure 7A, B and Supplementary figure 6 A-D)**. The immunoblot analysis of ATG8 revealed two distinct bands between 15 KDa to 10 KDa, *vi*z., free ATG8 and lipidated ATG8 (ATG8-PE), which varied in intensities in the samples analysed. Interestingly, the intensity of the ATG8-PE band was low in unprimed L-799 but higher in unprimed Suraj compared to their respective control seedlings under stress. Contrastingly, primed stressed L-799 showcased higher intensified bands of ATG8 and ATG8-PE compared to unprimed stressed seedlings. In contrast, in primed Suraj, the intensity of ATG8-PE was higher compared to its unprimed seedlings under stress **(Figure 7C)**. *Constitutively stressed 1* (*COST1*) negative regulator of autophagy showcased downregulated expression by 2.22 and 2.50-fold in unprimed seedlings of both L-799 and Suraj compared to respective controls under stress. Interestingly, priming caused a differential effect on *COST1* expression under drought stress in these varieties as the expression decreased (3.75-fold) in L-799 but was elevated (4.94-fold) compared to respective unprimed seedlings **(Figure 7B and Supplementary figure 7B).**

**Figure 7:**
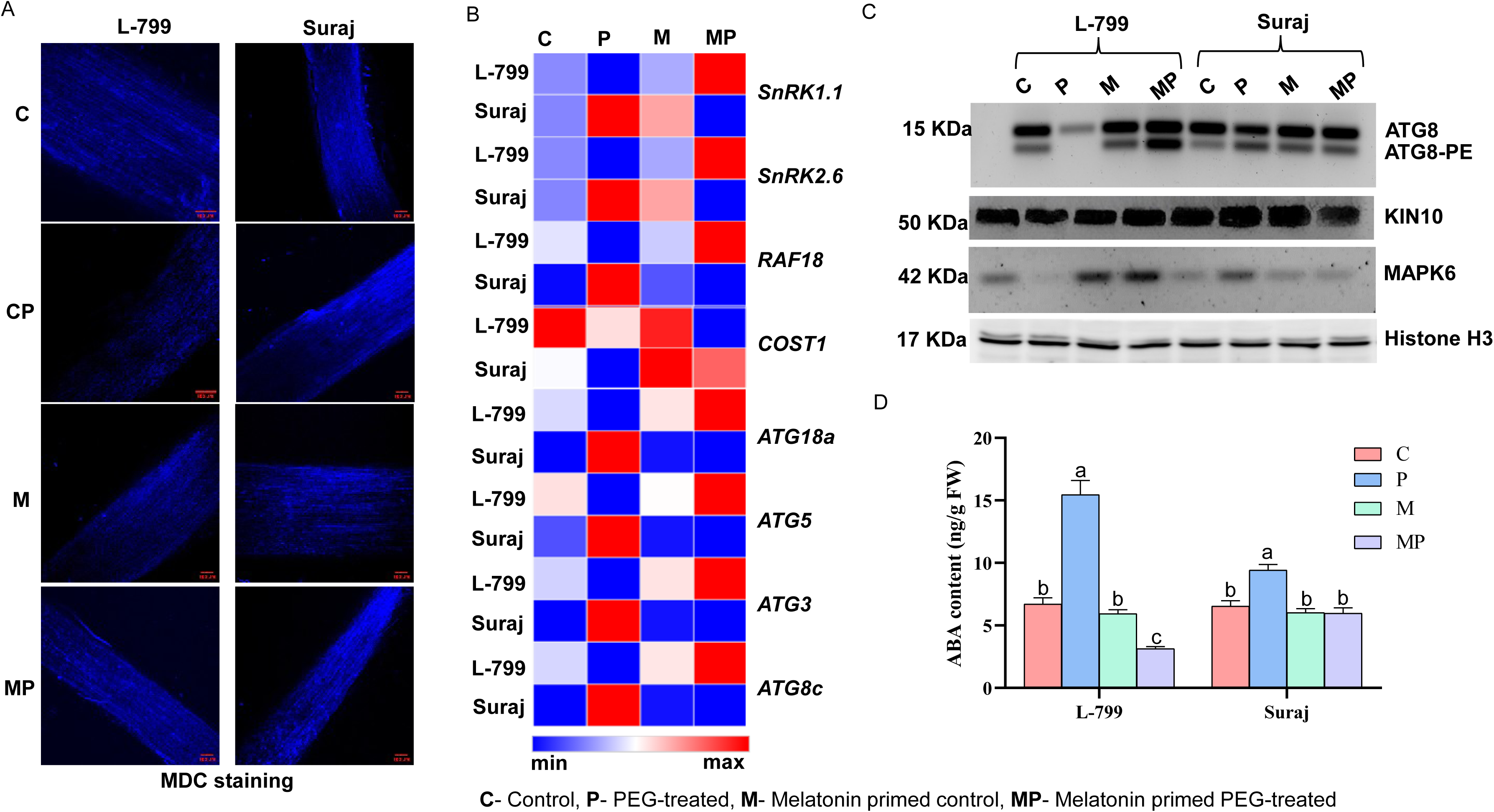
Effect of melatonin on autophagosome formation and expression of genes related to autophagy and ABA content under optimal and PEG-induced stress conditions in drought sensitive (L-799) and drought tolerant (Suraj) varieties of cotton (*Gossypium hirsutum* L.). (**A**) MDC staining, (**B**) Relative fold-change expression of genes related to autophagy *viz*., *SnRK1.1, SnRK2.1, RAF18, ATG18a, ATG5, ATG3, ATG8,* (**C**) Immunoblot analysis of ATG8, ATG8-PE, KIN10, MAPK6 and Histone H3, and (**D**) ABA content. The data represent mean values ± SE of 3 independent experiments, with 3 replicates per treatment in each experiment. Different alphabets within group represent significant differences among the treatments according to Duncan’s multiple range test (DMRT) at *p*-value ≤ 0.05.

ABA, a major phytohormone, is known for its role in imparting drought tolerance and autophagy in plants. The unprimed stressed seedlings of both varieties showed increased ABA content (2.30 times in L-799 and 1.44 times in Suraj) compared to respective controls. Melatonin priming decreased the intracellular ABA content by 4.92 and 1.57 times compared to unprimed seedlings under stress in both L-799 and Suraj, respectively (**Figure 7D**).

Sucrose non-fermenting1 related kinase (SnRKs) acts as a positive regulator of autophagy. The downregulated expressions of *SnRK2.6* (6.60-fold) and *SnRK1.1* (6.73-fold), along with the reduced intensity of KIN10 protein in unprimed stressed L-799 compared to controls, indicate reduced autophagy. Priming enhanced the expression of *SnRK2.6* (38.80-fold) and *SnRK1.1* (39.46-fold) and the protein intensity of KIN10 compared to unprimed seedlings of L-799 under stress. The unprimed seedlings of Suraj displayed significantly upregulated expression of *SnRK2.6* (4.17-fold), *SnRK1.1* (4.17-fold) and an intensified band of KIN10 protein compared to primed seedlings under stress (**Figure 7B, C Supplementary figure 6E and F)**.

MAP kinase (MAPK), another important cellular signalling molecule, can regulate several molecular pathways. Upon priming, L-799 exhibited significantly higher expression of *Rapidly Accelerated Fibrosarcoma 18* (*RAF18*) (196.-fold), a B4 MAPK and MAPK6 protein level compared to unprimed seedlings under stress conditions. Whereas, Suraj under stress exhibited higher levels of *RAF18* transcripts (19.28-fold) and MAPK6 protein in unprimed seedlings compared to primed ones (**Figure 7B, C and Supplementary figure 7A).**

### 3.10. Role of melatonin on cell viability and cell death

Water loss during drought stress coupled with increased osmotic stress compromises cell membrane integrity. Under drought stress, primed L-799 seedlings displayed lower intensity upon staining, with Evans blue indicating high cell viability compared to unprimed seedlings. However, Suraj exhibited no significant difference between primed and unprimed seedlings (**Figure 8A**). Genes such as bcl-2-associated athanogene 2 (*BAG*2), *BAG6*, metacaspase 9 (*MC9*), and suppressor of gamma response 1 (*SOG1*) play pivotal roles in cell death. Under PEG-induced drought stress, unprimed L-799 seedlings exhibited elevated expression of *BAG2* (1185.81-fold), *BAG6* (11.37-fold), *MC9* (10.38-fold), and *SOG1* (66.33-fold) genes, indicating higher cell death compared to the controls. Primed seedlings of L-799 showcased reduced transcript levels, *viz.*, *BAG2* (919.23-fold), *BAG6* (6.38-fold), *MC9* (5.16-fold), and *SOG1* (7.52-fold), compared to unprimed seedlings under drought. In the Suraj variety, although unprimed stressed seedlings exhibited higher expression of *BAG6* (2.01-fold) and *MC9* (1.65-fold) compared to controls, their expressions were lower in comparison to L-799. However, no significant change was observed in *BAG2* and *SOG1* expression in unprimed and primed seedlings of Suraj under drought stress conditions **(Figure 8B and Supplementary figure 7C-F)**.

**Figure 8:**
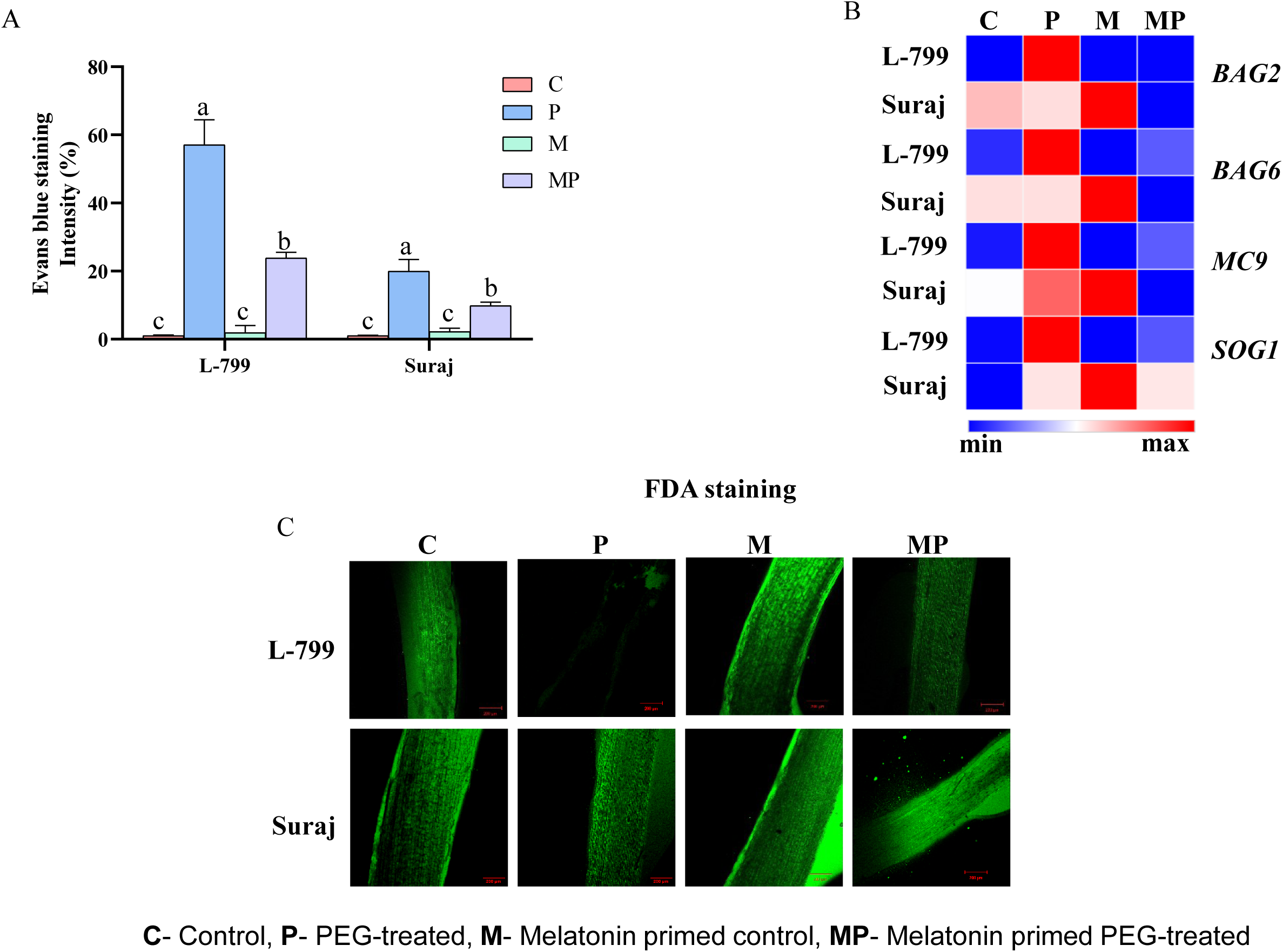
Effect of melatonin priming on cell viability, necrosis and expression of related genes under optimal and PEG-induced stress conditions in drought-sensitive (L-799) and drought-tolerant (Suraj) varieties of cotton (*Gossypium hirsutum* L.). (**A**) Cell death assessed by Evans blue staining, (**B**) Relative fold change expression of necrosis related genes *BAG2, BAG6, MC9* and *SOG1,* and (**C**) Detection of viable cells by FDA staining. The data represent mean values ± SE of 3 independent experiments, with 3 replicates per treatment in each experiment. Different alphabets within the group represent significant differences among the treatments according to Duncan’s multiple range test (DMRT) at *p*-value ≤ 0.05.

**Figure 9:**
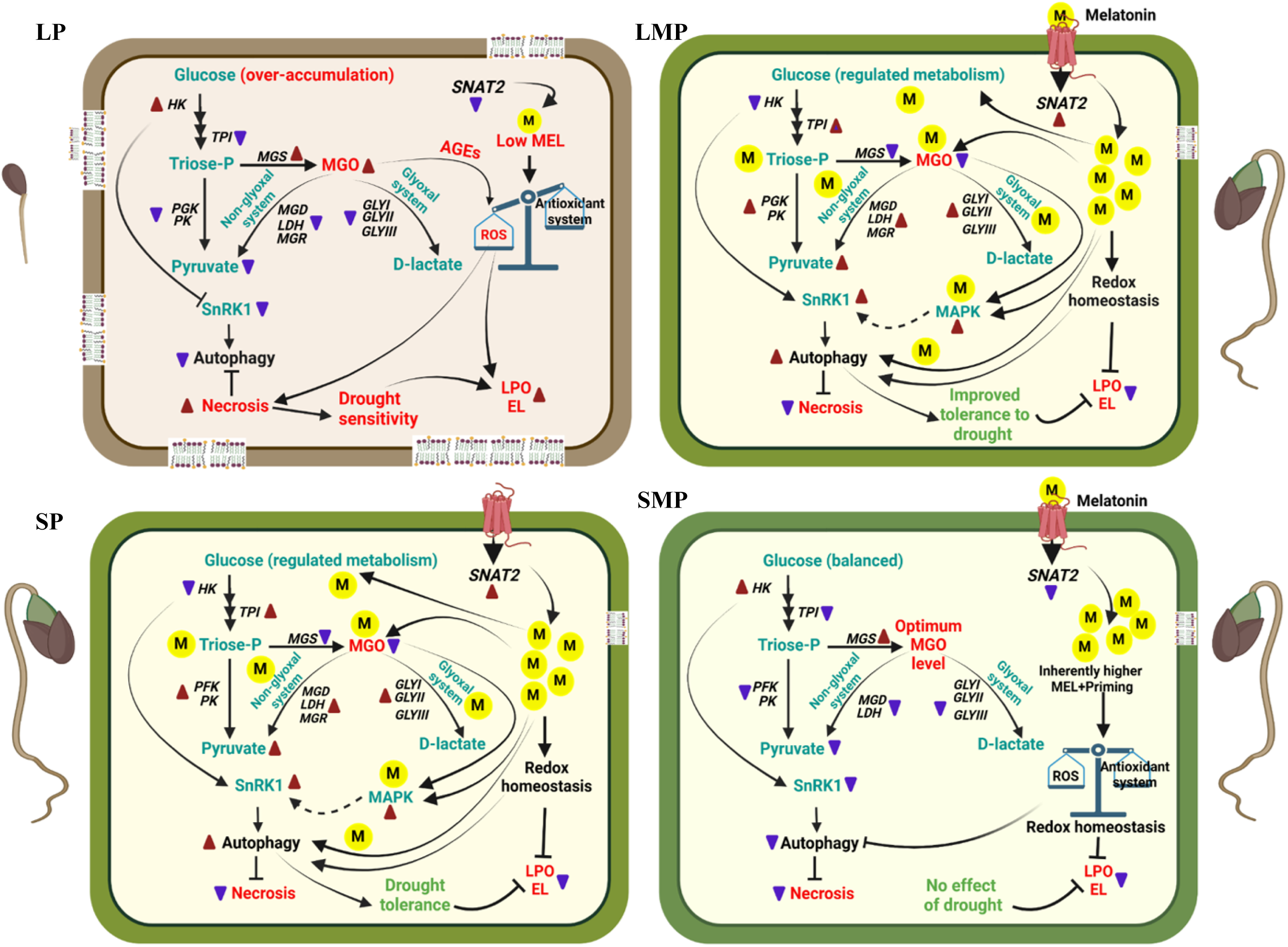
Diagrammatic illustration of melatonin-mediated differential regulation of MGO-homeostasis and autophagy under PEG-induced drought stress during seed germination in drought-sensitive (L-799) and drought-tolerant (Suraj) varieties. Lower melatonin and higher accumulation of MGO in unprimed L-799 leads to necrosis under drought stress (**LP**). Melatonin aids in better conversion of glucose to pyruvate and improved glyoxalase and non-glyoxalase system thus helping in maintenance of intracellular MGO homeostasis. The maintained MGO, by acting as signaling molecule, upregulates the MAPK expression, which in-turn activates SnRK1 to induce autophagy and consequently improves drought tolerance in L-799 (**LMP**). The higher endogenous melatonin content, better MGO homeostasis, redox regulation, and autophagy under drought in the drought-tolerant variety could be attributed to its drought-tolerant nature (**SP**). Priming allowed maintenance of normal metabolic functions without the need for additional stress responses in Suraj due to its drought tolerant nature. It balanced the cellular stability, without requiring further metabolic adjustments, such as autophagy, antioxidant activity, ROS, intracellular sugar level, showcasing no further beneficial effect of priming in this variety (**SMP**). LP, Unprimed L-799 seedlings grown under PEG induced drought stress. LMP, Melatonin primed L-799 seedlings grown under PEG induced drought stress. SP, Unprimed Suraj seedlings grown under PEG induced drought stress. SMP, Melatonin primed Suraj seedlings grown under PEG induced drought stress.

Cell viability, assessed using FDA staining, distinguishes live and dead cells by detecting intercellular esterase activity. In L-799, primed seedlings displayed higher staining (indicating more viable cells) than unprimed seedlings with lower staining (indicating less viable cells) under stress. In Suraj, both primed and unprimed seedlings showed higher staining intensity under stress **(Figure 8C)**.

## 4. Discussion

MGO is spontaneously produced and over accumulates during drought and other abiotic stresses and thus poses a threat to cellular health (Askari-Khorasgani and Pessarakli 2019). Several molecules have emerged as priming agents to mitigate the detrimental impact of drought stress (Paparella et al. 2015). Melatonin is one such molecule which has been reported to improve germination and seedling growth under stress conditions (Ahmad et al. 2021; Awan et al. 2023). In the present study, melatonin priming at 25 μM was effective in improving the seedling phenotypes of L-799 as well as relative water content whereas no significant differences were noticed in primed Suraj as compared to unprimed seedlings under stress (**Figure 1D**). The differences in endogenous melatonin content in drought distinguished varieties could be a reason for the differential responses to melatonin priming under PEG-induced drought stress conditions as suggested by (Supriya et al. 2022).

### 4.1. Melatonin priming raised endogenous melatonin levels and regulated intracellular ROS and AGEs under PEG-induced drought stress

Intracellular melatonin plays a crucial role in mitigating the stress-induced detrimental effects on plants (Zhang et al. 2014), while exogenous application is reported to improve the endogenous content, especially under drought stress conditions (Chen et al. 2020; Guo et al. 2022; Supriya et al. 2022). The PMTR1 mediated signaling is known to regulate seed germination and seedling growth (Yin et al. 2022), leaf senescence (Bai et al. 2022), stomatal closure and circadian stomatal rhythm *via* MAPK (Lee and Back 2016; Yang et al. 2021) under osmotic and drought stress (Wang et al. 2021). Notably, in L-799, elevated melatonin levels correlated with upregulated expression of the *PMTR1* and *SNAT2* genes in primed seedlings than unprimed under stress, indicating the efficacy of exogenous priming to elevate the endogenous content thus improving the stress tolerance. Elevated endogenous melatonin content, *PMTR1* and *SNAT2* expression, coupled with better stress tolerance in unprimed conditions compared to controls, can be explained by Suraj’s inherent drought-tolerant nature. Contrastingly, despite elevated *PMTR1*, insignificant changes in melatonin content and a decrease in *SNAT2* expression was observed in primed Suraj compared to unprimed under stress. This discrepancy explains that exogenous priming primarily enhances *PMTR1* expression, while the elevated melatonin content trigger feedback inhibition of the *SNAT* gene, thereby regulating endogenous melatonin content in primed Suraj, as reported earlier by Shreya et al. (2022) **(Figure 2A and B, Supplementary figure 2A and B)**.

Drought disrupts cellular redox potential by elevating the ROS production, ultimately disorders cellular antioxidant harmony and increases lipid peroxidation and EL (Cruz de Carvalho 2008). Moreover, drought also disrupts glycolytic flux, leading to the accumulation of triose phosphate intermediates which are converted to MGO (Hossen et al. 2022). Furthermore, Hussain et al. (2018) and Hasan et al. (2020) reported that drought-induced MGO production triggers ROS accumulation through AGEs formation and can also be another reason for higher production of H_2_O_2_, MDA and EL leading to increased cellular damage in both the varieties under stress. Melatonin attenuates the ROS generation and AGEs formation and thus exhibits anti-aging properties (Sehirli et al. 2021). Decreased AGE-inhibition potential under drought compared to controls in L-799 signifies the increase in AGEs formation and consequential harmful impact of the stress. However, priming-mediated enhanced AGE-inhibition activity in L-799 justifies the protective effect of melatonin against AGE formation, similar to the result of (Takabe et al 2016). Contrastingly, the higher AGE-inhibition activity in unprimed Suraj under stress could be due to elevated endogenous melatonin content and its inherent tolerant nature. Primed seedlings of both the varieties exhibited significantly alleviated ROS accumulation (lesser staining of DAB, NBT and H_2_O_2_ content) and therefore, decreased the EL levels and MDA under PEG-induced drought stress conditions, justifying the beneficial effect of melatonin in regulating redox homeostasis (**Figure 3A-E and 4B**).

### 4.2. Melatonin maintained intracellular MGO-homeostasis by improving glyoxalase, non-glyoxalase system, glutathione pool and glycolysis

The interplay between disrupted redox potential and increased MGO levels creates a vicious cycle of oxidative stress, compromising seedling germination and growth under drought conditions. To mitigate the overaccumulation of this potential cytotoxic compound, plants initiate glyoxalase systems to detoxify and convert MGO into less harmful compounds (Talaat and Todorova 2022). Seedlings of unprimed L-799 exhibited significantly higher intracellular MGO with upregulated MGO-biosynthesis genes (*MGS*1 and *SSAO1*), which correlated with the decreased enzymatic activities (GLY I, II, III) and transcript expressions of *GLY I*, *II*, *III*, *MGR1*, *MGD1* and *LDH5,* showing the negative effect of drought stress in L-799 (drought-sensitive variety). Interestingly, melatonin priming notably elevated activities (GLY I, II, III) and expressions of *GLY I*, *II*, *III*, *MGR1*, *MGD1* and *LDH5* in L-799, justifying the beneficial effect of melatonin on the glyoxalase system to decrease intracellular MGO levels compared to unprimed under PEG-induced stress. Nevertheless, the reduced MGO levels coupled with decreased expression of *MGS1* and *SSAO1* along with augmented activities and expressions of GLY I, II, and III in unprimed Suraj under stress conditions, suggest its inherent potential to withstand stress (**Figure 4A, C, 5A-C, E and Supplementary figure 2C, D and 3A-F**). Our findings are in agreement with other reports wherein melatonin application has been shown to improve the plant’s glyoxalase and non-glyoxalase antioxidant systems and thus efficiently scavenged excess MGO and maintained cellular homeostasis (Hussain et al. 2018; Kaya et al. 2023).

GSH is a fundamental player in maintaining cellular redox homeostasis by scavenging ROS and MGO under stress (de Bari et al. 2020). Yadav et al. (2005) reported that an increase in MGO levels correlated with decrease in the GSH levels under stress. In this study, primed seedlings with significantly increased GSH and GSH/GSSH ratio was associated with enhanced *GR1* expression compared to unprimed seedlings under drought stress in L-799, justifying melatonin’s protective role against MGO through the maintenance of the GSH pool, consistent with the findings of Kaur and Bhatla (2016). Contrastingly, non-significant differences in GSH level and *GR1* expression between primed and unprimed seedlings of Suraj under drought stress suggest no additional beneficial effect of melatonin in the tolerant variety as reported by Supriya et al. (2022) (**Figure 5D, E and Supplementary figure 4A-D**).

Abiotic stress, particularly drought stress mediated injury results in sugar accumulation (Kaur et al. 2021). Sugar metabolism is directly linked to MGO production, as reported by Borysiuk et al. (2018). Elevated sugar levels enhance glycolysis, resulting in increased production of MGO. This concomitantly diminishes the capacity of glycolytic enzymes such as TPI and downstream of triphosphates to manage the augmented glucose flux. Melatonin-primed L-799 displayed a significant decrease in the glucose content (2.24 times) along with elevated pyruvate content (6.58 times) compared to unprimed seedlings under stress. Melatonin priming lowered the expression of glycolysis upstream gene *HK3* but upregulated *TPI1*, *PGK5* and *PK1* compared to unprimed seedlings in L-799 under stress conditions. Conversely, in Suraj, the levels of glucose, pyruvate and the expression of glycolysis genes exhibited opposite trend compared to L-799 in primed and unprimed seedlings under stress conditions. Thus, it can be suggested that melatonin aids in metabolizing accumulated sugars to pyruvate by enhancing glycolysis, especially downstream genes, therefore reducing MGO formation. The increased *TPI1* expression levels due to melatonin priming under stress are supported by Sharma et al. (2012) findings, where increased expression of TPI regulates the swift equilibrium between DHAP and GAP, thereby reducing the toxic levels of MGO (**Figure 6A, B and Supplementary figure 5A-F**).

### 4.3. Melatonin priming-induced MGO homeosis was associated with enhanced autophagosome formation

Autophagy plays a crucial role in plant survival and stress response (Su et al. 2020). Cui et al. (2018) found that melatonin can ameliorate seed germination by improving energy production and activating a metabolic cascade related to autophagy, protein degradation, and other processes under PEG stress. Higher MDC-stained (autophagosome) bodies correlated with upregulated expression of *ATG*s and (ATG8-PE:ATG8) in primed L-799, pointing towards a positive correlation between melatonin priming and autophagy induction under stress conditions, similar to the results of Supriya et al. 2022). Cellular energy status and autophagy are inter-dependent attributes, as energy deprivation incites autophagy to activate (Feng et al. 2024), and simultaneously autophagy also improves cellular energy status by reusing the non-functional macromolecules (He 2022). The decreased ATP content in L-799 primed seedlings compared to controls under stress, justifies its energy-deprived state which correlated to higher autophagosome content. But, the decreased ATP content in unprimed seedlings compared to primed seedlings can be the reason of weak autophagy signal as depicted by lesser autophagosome formation and *ATGs* expression in L-799, as already reported by (Supriya et al. 2024). Lower ATP content correlated to higher autophagosome and ATGs expressions in unprimed Suraj, justifying the incitation of autophagy under energy deprivation conditions. Surprisingly, no change in energy (ATP) content along with lesser expression levels of *ATG*s in primed stressed compared to unprimed control substantiates the intracellular balanced state of Suraj. Autophagy known to prevent MGO-induced apoptosis through activation of AMPK (Park et al. 2020). Furthermore, higher expressions of *SnRK2.6*, *SnRK1.1* and KIN10 which are positive regulators of autophagy (Yang et al. 2023), along with downregulated *COST1*, a negative regulator of autophagy (Bao et al. 2020), in primed stressed plants of L-799 further supports the better autophagy induction. Interestingly, higher *ATG*s and ATG-PE:ATG8 expressions, MDC-stained bodies, upregulated *SnRK2.6*, *SnRK1.1* and KIN10 expressions along with alleviated expressions of *COST1* in unprimed Suraj, suggests the improved autophagy which might be due to higher endogenous melatonin content. Moreover, decreased autophagy induction as reflected from lesser autophagosome and *ATG*s expression in primed Suraj compared to unprimed shows that exogenous priming is unable to mediate further improvement in stress responses in drought tolerant variety (**Figure 6C, 7A-C and Supplementary figure 6A-F, 7B**).

### 4.4. MGO signaling might activate autophagy *via* MAPK, independent ABA

Abscisic acid (ABA) plays dual roles as a negative regulator of seed germination and positive regulator of autophagy (Sirko et al. 2021). Under drought conditions, increased levels of ABA compared to controls, shows the negative effect of stress on germination in unprimed L-799. Conversely, melatonin priming mitigated this effect by reducing ABA levels, thereby promoting seedling growth in L-799 compared to unprimed under stress conditions. Despite melatonin’s antagonistic relationship with ABA, it paradoxically induced autophagy, presenting a scientific conundrum. In the recent study (Supriya et al. 2024), we reported that melatonin induced autophagy independent of ABA through the activation of MAPK. Notably, previous studies in animals demonstrated that the interaction of AGEs with plasma membrane receptors known as receptor for advanced glycation end-products (RAGEs), activated MAPK cascades in response to defense (Lee et al. 2014; Jeong and Lee 2021). Here, primed seedlings displayed a significant increase in the expression of *RAF18* and MAPK6 which correlated with higher transcript levels of *SnRK2.6*, *SnRK1.1* and KIN10 in L-799 compared to unprimed under stress, thus indicating a positive impact of melatonin on autophagy under stress similar to the findings of (Supriya et al. 2024). Honig et al. (2012) reported that overexpression of Atg8-interacting proteins (ATI1 and ATI2) stimulated seed germination in the presence of ABA, justifying the elevated levels of ABA along with autophagy in unprimed Suraj under stress. Thus, this study may explain that melatonin can potentially reduce drought-induced MGO levels, thereby acting as a signaling molecule that activates MAPK and promotes autophagy, which is beneficial for seed germination (**Figure 7B-D, Supplementary figure 6E, F and 7A**).

### 4.5. Melatonin regulated drought-induced cell death during MGO homeostasis

MGO can induce both apoptosis and autophagy, but the determining conditions between the two pathways remain unclear (Park et al. 2020). MGO, precursor for AGEs formation induces intracellular damage by increasing ROS levels and mitochondrial damage, leading to apoptosis (Park *et al*., 2020). Parallelly, existing research on animals and humans reports that autophagy plays a crucial role in mitigating MGO-induced apoptosis (Park et al. 2020), which has not yet been addressed in plant systems. Under drought, increased MGO levels with higher cell death and elevated expression of *BAG2*, *BAG6*, *MC9*, and *SOG1* in unprimed seedlings compared to controls in both the varieties shows that the drought induced cell-death could be a result of elevated intracellular MGO content. Similarly, significantly downregulated MGO content with reduced expression of necrosis related genes and decreased cell death in primed L-799 substantiates the beneficial role of melatonin against cell death under drought, further confirmed by higher autophagy. Interestingly, unprimed Suraj under stress exhibited elevated expression of necrosis genes (*BAG6* and *MC9*), and higher cell death which correlated with high ROS levels and *ATGs* expression. This suggests a mechanism in which both cell death and autophagy help in balancing cellular status during stress, highlighting Suraj’s inherent drought tolerance. These findings align with studies showing ROS can trigger autophagy (Yoshimoto et al. 2009; Wang et al. 2015) and has a dual role, supporting survival through autophagy at optimal levels but leading to necrotic cell death when produced at excess levels during prolonged exposure (Sadhu et al. 2019). In Suraj, melatonin priming did not significantly alter MGO content or the expression of *BAG2* and *SOG1*; however, it reduced *BAG6*, *MC9* expression and ROS levels and associated cell death without compromising its inherent tolerant nature (**Figure 8 and Supplementary figure 7C-F**).

## 5. Summary

The present study is a significant step towards unravelling the molecular and physiological intricacies of melatonin-mediated regulation of intracellular methylglyoxal during seed germination under PEG induced drought stress. Melatonin priming increased the endogenous melatonin content, thereby enhancing the glyoxalase and non-glyoxalase system, leading to reduced MGO accumulation in drought-sensitive (L-799) variety under stress. Despite reduced ABA content, melatonin-mediated MGO homeostasis might have induced autophagy by activating *SnRK2* and KIN10 *via* elevated MAPK6 levels. Conversely, in the drought-tolerant (Suraj) variety, priming did not cause marked changes in these attributes compared to unprimed seedlings under stress, which can be explained by higher endogenous melatonin content and inherent drought tolerance (**Figure 9**). Furthermore, this study is the first to report the autophagy-inducing effect of MGO under drought stress at optimal levels in upland cotton. Further studies using transgenic approaches would unveil additional intricacies involved in plant responses to stress, offering insights into potential strategies for enhancing plant resilience to changing environmental conditions.

## Author contributions

GP, DD and LS contributed to the conceptualization and experiment design. DD conducted the experiments, data collection, graph preparation, analysis, interpreted the data and wrote the manuscript. LS performed a few experiments, interpreted the data, corrected the manuscript, and helped with the pictorial representation of the summary. AK performed a few experiments. GP analysed and interpreted the results and corrected the manuscript. All authors contributed to the work and edited the manuscript.

## Acknowledgments

We would like to thank Dr. B. Sree Lakshmi, Principal Scientist and Head Cotton Section, Regional Agricultural Research Station, Acharya N. G. Ranga Agricultural University, Guntur, Andhra Pradesh, India for providing seeds of L-799 variety, and the Director, ICAR-Central Institute for Cotton Research (CICR), Nagpur, India for seeds of Suraj variety used in the research work. We would also like to thank Ayon Chatterjee and Arpan Chatterjee, Dept. of Biochemistry, University of Hyderabad, for their help in carrying out the experiments involving HPLC and western blot.

## Data availability statement

Data sharing is not applicable to this article as all new created data is already contained within this article.

The data that supports the findings of this study are available in the supplementary material of this article

