## Supplemental Figure 1, Supplemental Figure 2, Supplemental Figure 3, Supplemental Figure 4, Supplemental Figure 5, Supplemental Figure 6, Supplemental for "Melatonin-mediated methylglyoxal homeostasis and regulation of autophagy during seed germination under PEG-induced drought stress in upland cotton"

### Slide 1
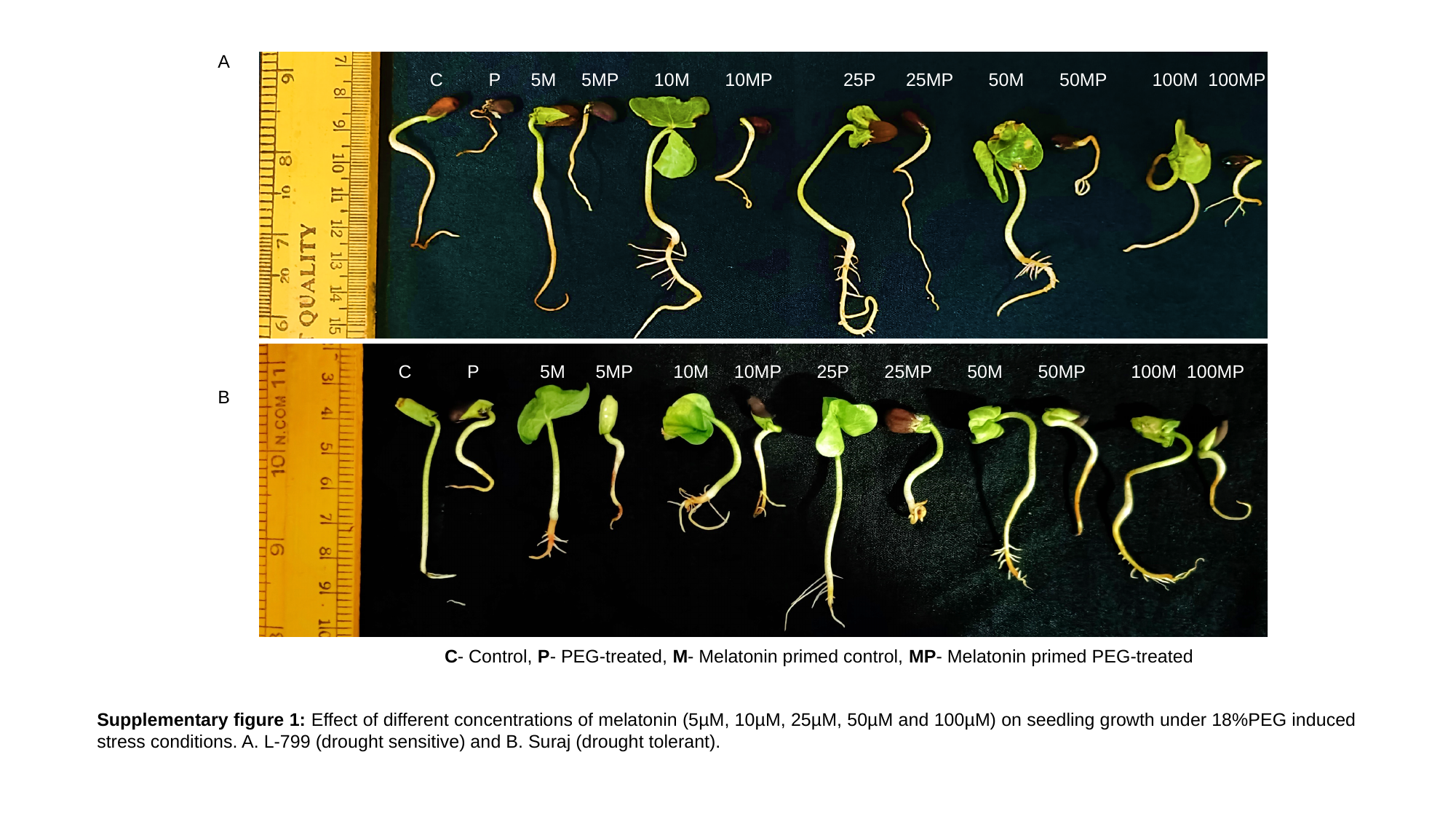

A
 C P 5M 5MP 10M 10MP 25P 25MP 50M 50MP 100M 100MP
 C P 5M 5MP 10M 10MP 25P 25MP 50M 50MP 100M 100MP
 C P 5M 5MP 10M 10MP 25P 25MP 50M 50MP 100M 100MP
B
C- Control, P- PEG-treated, M- Melatonin primed control, MP- Melatonin primed PEG-treated
Supplementary figure 1: Effect of different concentrations of melatonin (5µM, 10µM, 25µM, 50µM and 100µM) on seedling growth under 18%PEG induced stress conditions. A. L-799 (drought sensitive) and B. Suraj (drought tolerant).

### Slide 2
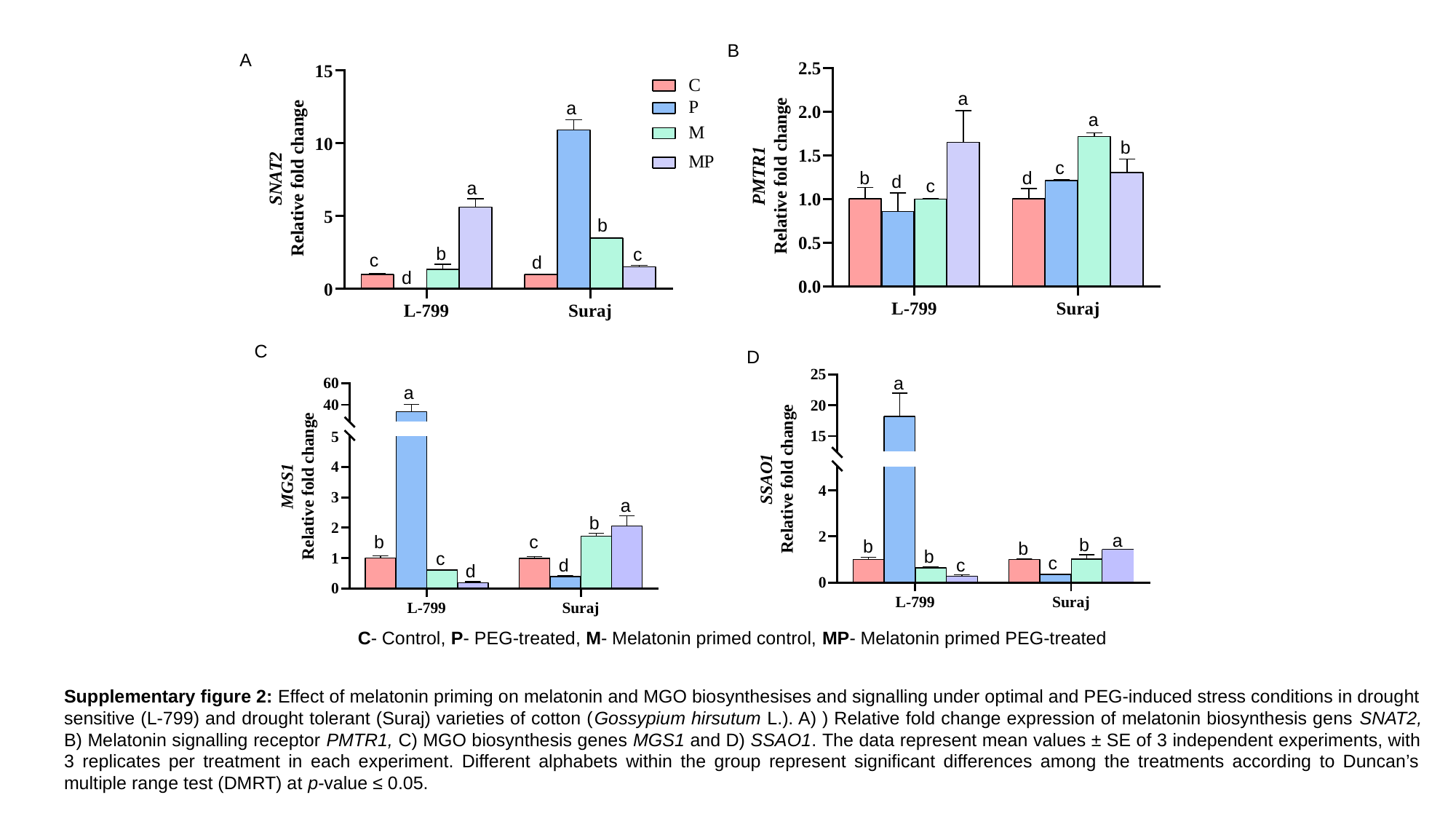

B
A
a
a
a
b
c
d
b
d
c
a
b
b
c
c
d
d
C
D
a
a
a
b
b
c
c
d
d
a
b
b
b
b
c
c
C- Control, P- PEG-treated, M- Melatonin primed control, MP- Melatonin primed PEG-treated
Supplementary figure 2: Effect of melatonin priming on melatonin and MGO biosynthesises and signalling under optimal and PEG-induced stress conditions in drought sensitive (L-799) and drought tolerant (Suraj) varieties of cotton (Gossypium hirsutum L.). A) ) Relative fold change expression of melatonin biosynthesis gens SNAT2, B) Melatonin signalling receptor PMTR1, C) MGO biosynthesis genes MGS1 and D) SSAO1. The data represent mean values ± SE of 3 independent experiments, with 3 replicates per treatment in each experiment. Different alphabets within the group represent significant differences among the treatments according to Duncan’s multiple range test (DMRT) at p-value ≤ 0.05.

### Slide 3
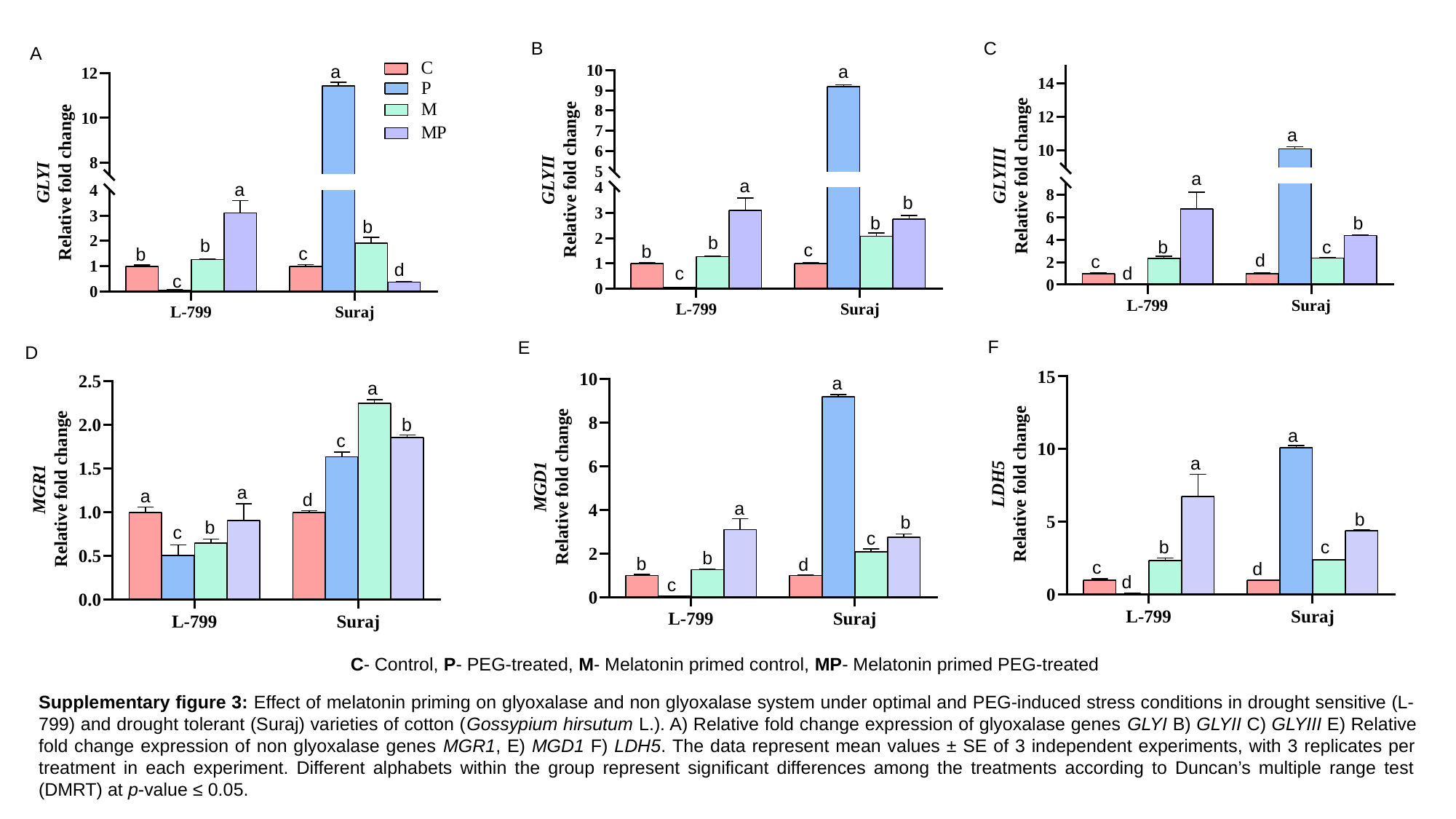

B
C
A
a
a
a
a
a
a
b
b
b
b
b
b
c
b
c
b
c
b
d
c
d
d
c
c
F
E
D
a
a
b
c
b
c
d
d
a
a
b
c
b
b
d
c
a
b
c
a
a
d
b
c
C- Control, P- PEG-treated, M- Melatonin primed control, MP- Melatonin primed PEG-treated
Supplementary figure 3: Effect of melatonin priming on glyoxalase and non glyoxalase system under optimal and PEG-induced stress conditions in drought sensitive (L-799) and drought tolerant (Suraj) varieties of cotton (Gossypium hirsutum L.). A) Relative fold change expression of glyoxalase genes GLYI B) GLYII C) GLYIII E) Relative fold change expression of non glyoxalase genes MGR1, E) MGD1 F) LDH5. The data represent mean values ± SE of 3 independent experiments, with 3 replicates per treatment in each experiment. Different alphabets within the group represent significant differences among the treatments according to Duncan’s multiple range test (DMRT) at p-value ≤ 0.05.
b
b
a

### Slide 4
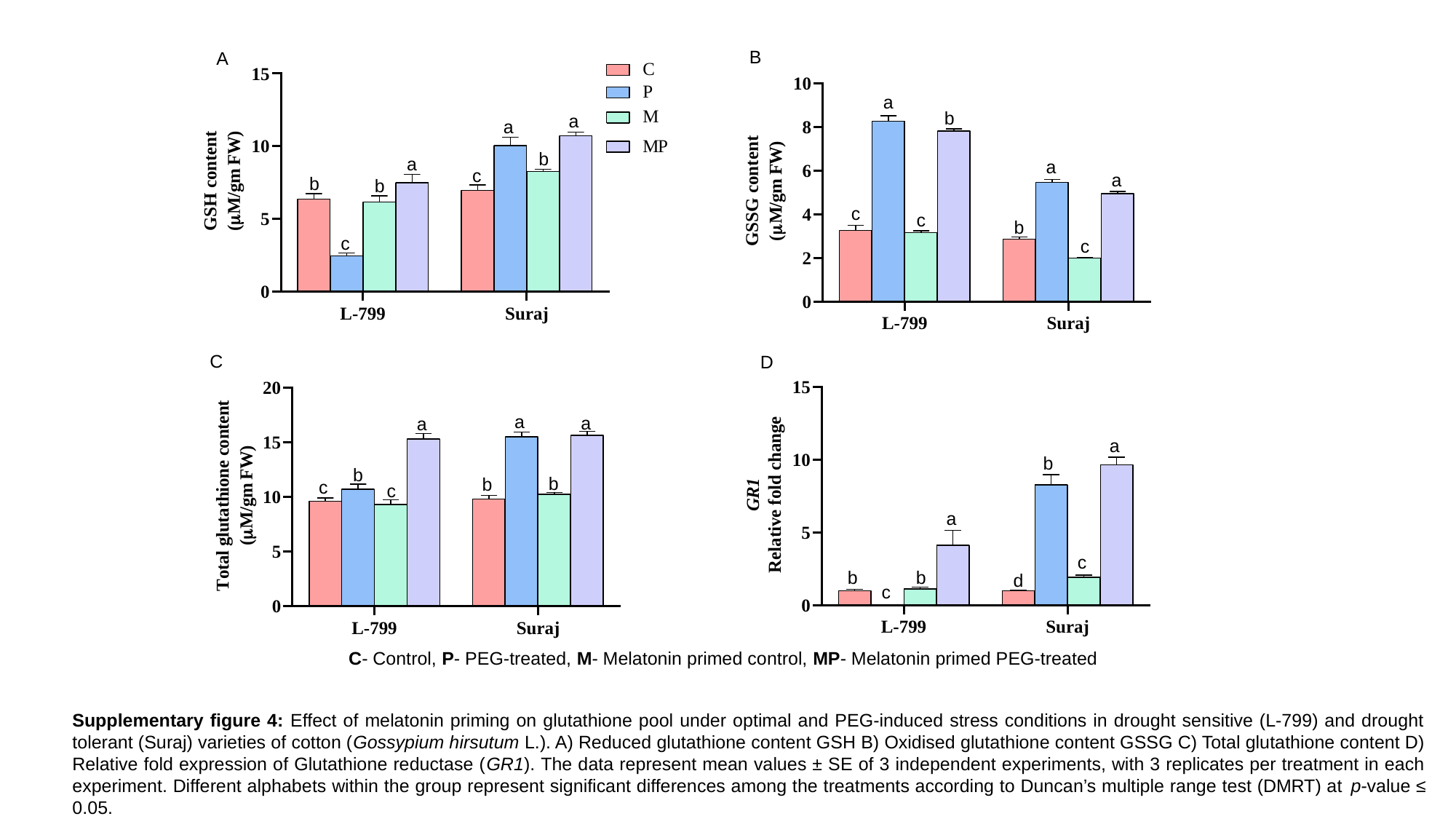

B
a
b
a
a
c
c
b
c
A
a
a
b
a
c
b
b
c
C
D
a
a
a
b
b
b
c
c
a
b
a
c
b
b
d
c
C- Control, P- PEG-treated, M- Melatonin primed control, MP- Melatonin primed PEG-treated
Supplementary figure 4: Effect of melatonin priming on glutathione pool under optimal and PEG-induced stress conditions in drought sensitive (L-799) and drought tolerant (Suraj) varieties of cotton (Gossypium hirsutum L.). A) Reduced glutathione content GSH B) Oxidised glutathione content GSSG C) Total glutathione content D) Relative fold expression of Glutathione reductase (GR1). The data represent mean values ± SE of 3 independent experiments, with 3 replicates per treatment in each experiment. Different alphabets within the group represent significant differences among the treatments according to Duncan’s multiple range test (DMRT) at p-value ≤ 0.05.

### Slide 5
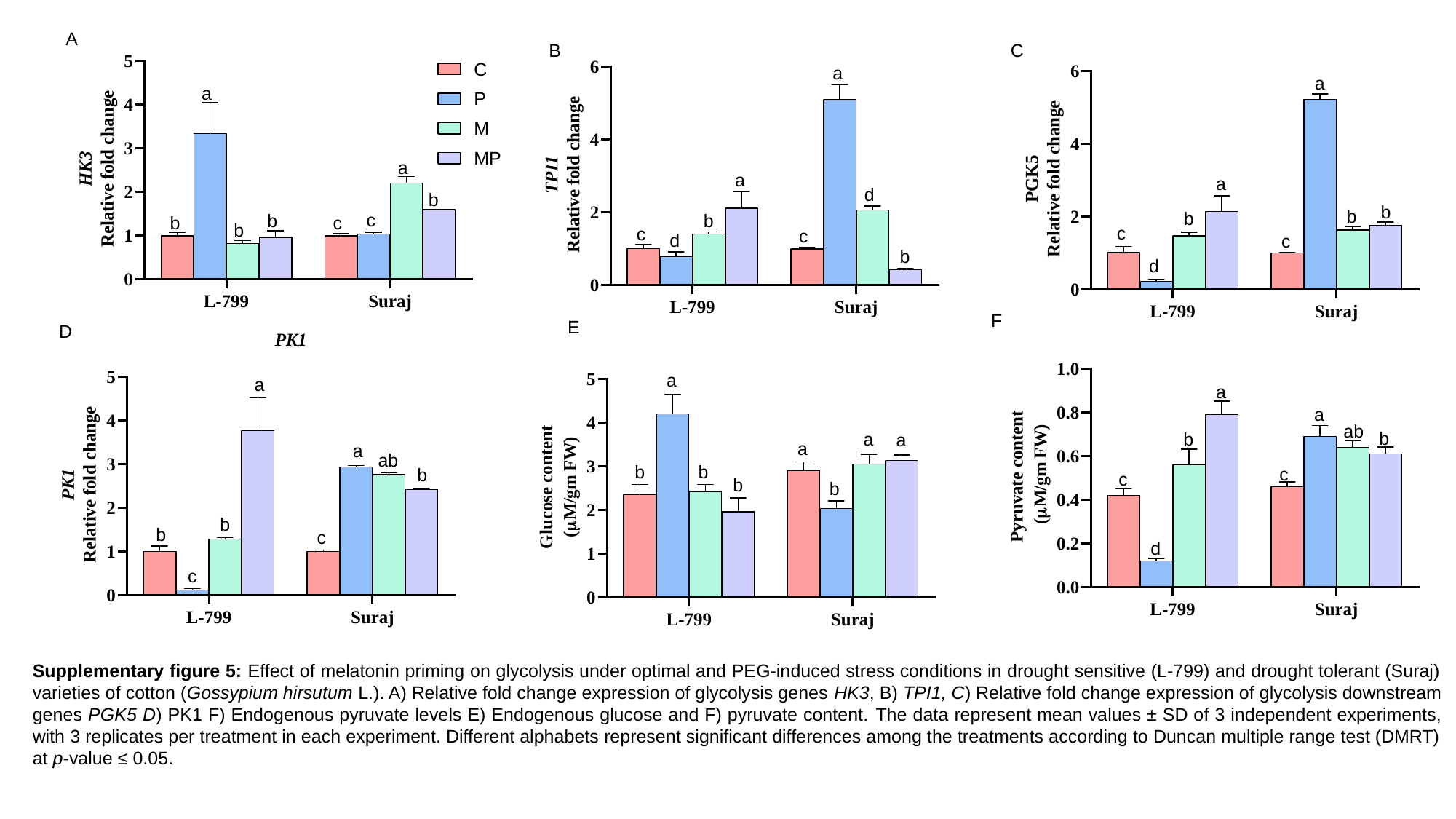

A
C
B
a
a
d
b
c
c
d
b
a
a
b
b
c
d
b
c
a
a
b
c
b
b
c
b
a
a
b
b
c
c
ab
b
F
E
D
a
a
a
a
b
b
b
b
a
a
ab
b
b
c
c
d
Supplementary figure 5: Effect of melatonin priming on glycolysis under optimal and PEG-induced stress conditions in drought sensitive (L-799) and drought tolerant (Suraj) varieties of cotton (Gossypium hirsutum L.). A) Relative fold change expression of glycolysis genes HK3, B) TPI1, C) Relative fold change expression of glycolysis downstream genes PGK5 D) PK1 F) Endogenous pyruvate levels E) Endogenous glucose and F) pyruvate content. The data represent mean values ± SD of 3 independent experiments, with 3 replicates per treatment in each experiment. Different alphabets represent significant differences among the treatments according to Duncan multiple range test (DMRT) at p-value ≤ 0.05.

### Slide 6
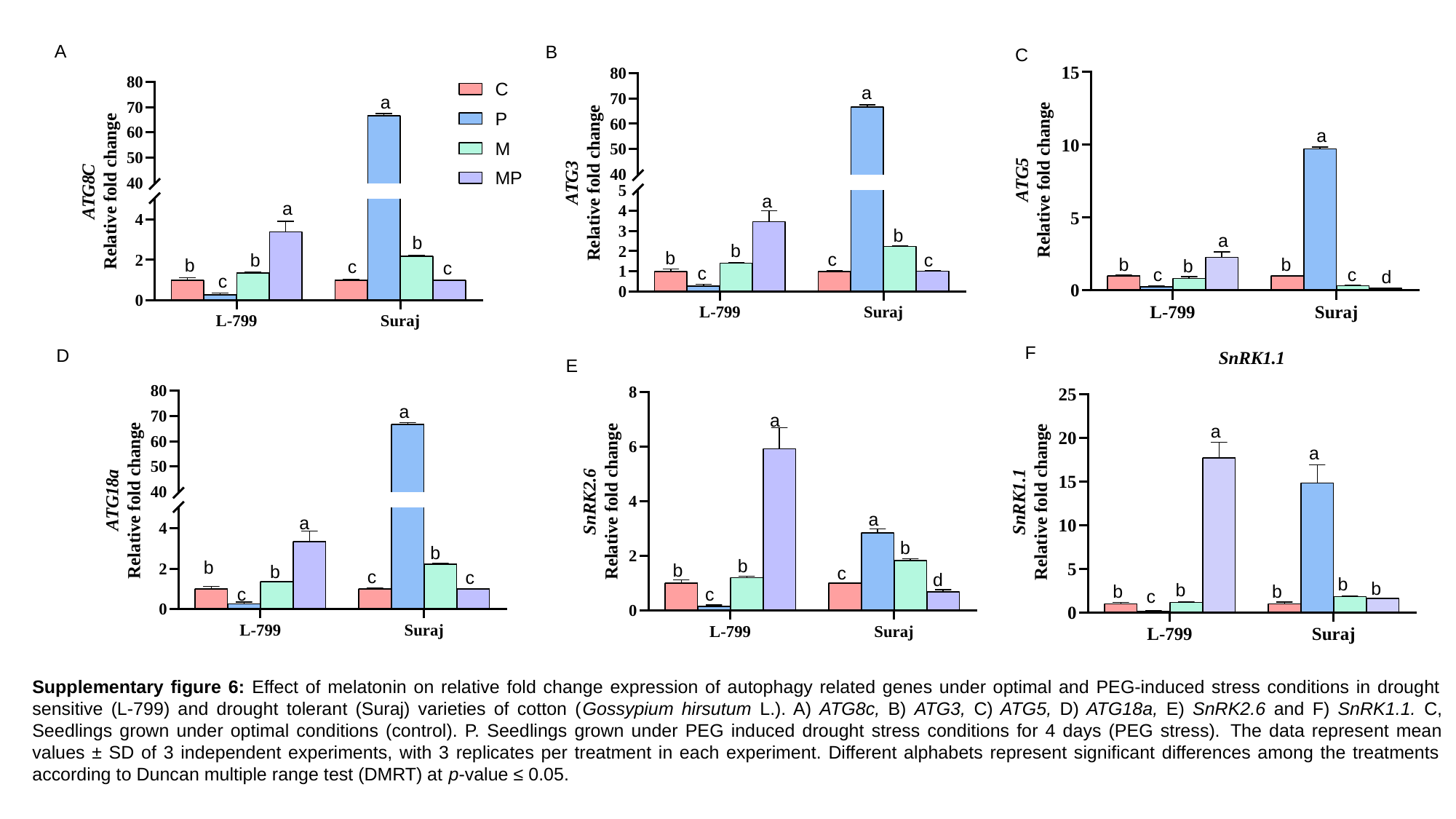

A
B
a
a
b
b
b
c
c
c
C
a
a
b
b
b
c
c
d
a
a
b
b
b
c
c
c
F
a
a
b
b
b
b
b
c
D
E
a
a
b
b
b
c
d
c
a
a
b
b
b
c
c
c
Supplementary figure 6: Effect of melatonin on relative fold change expression of autophagy related genes under optimal and PEG-induced stress conditions in drought sensitive (L-799) and drought tolerant (Suraj) varieties of cotton (Gossypium hirsutum L.). A) ATG8c, B) ATG3, C) ATG5, D) ATG18a, E) SnRK2.6 and F) SnRK1.1. C, Seedlings grown under optimal conditions (control). P. Seedlings grown under PEG induced drought stress conditions for 4 days (PEG stress). The data represent mean values ± SD of 3 independent experiments, with 3 replicates per treatment in each experiment. Different alphabets represent significant differences among the treatments according to Duncan multiple range test (DMRT) at p-value ≤ 0.05.

### Slide 7
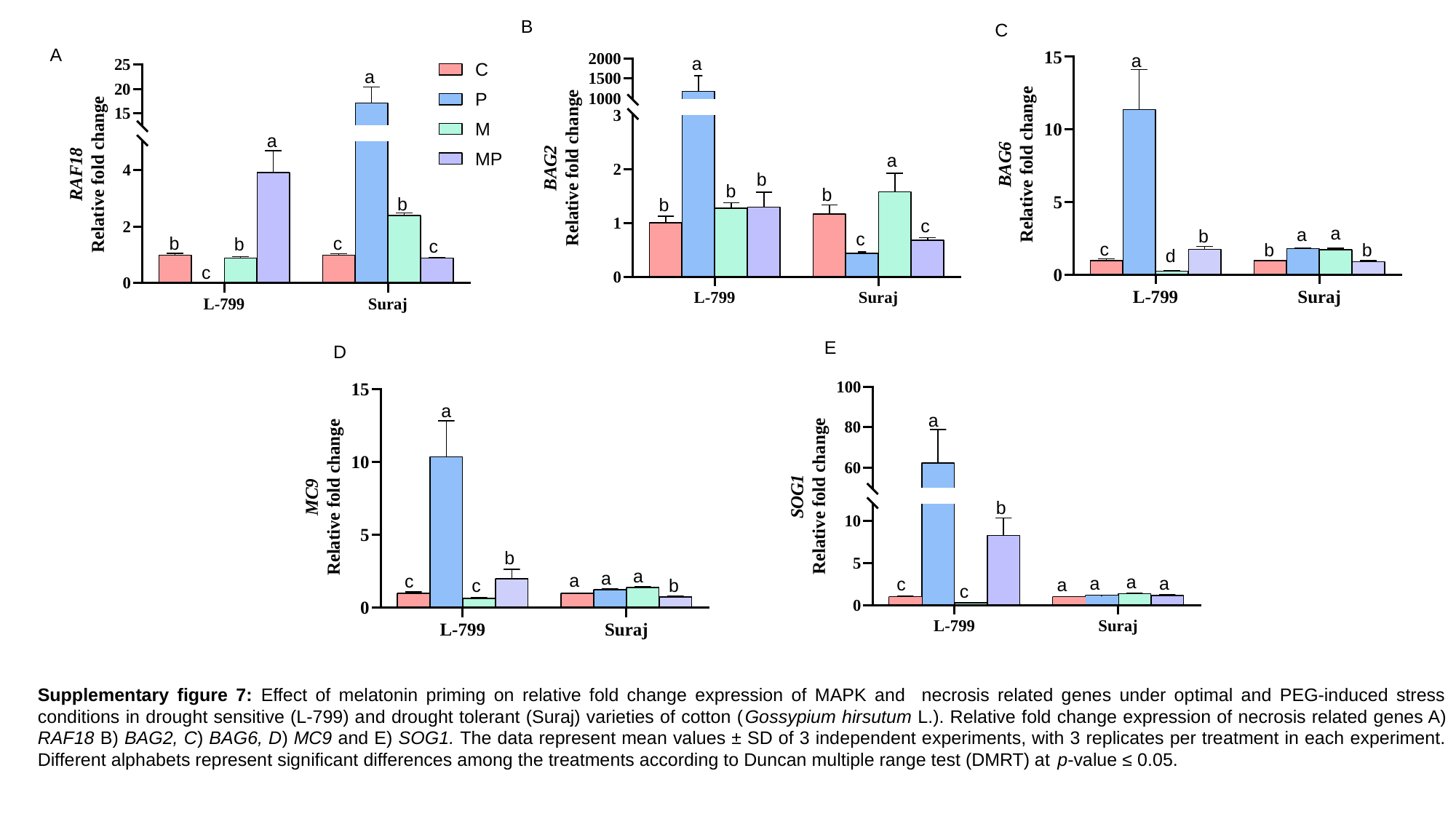

B
C
a
a
a
b
c
b
b
d
a
a
b
b
b
b
c
c
A
a
a
b
b
c
b
c
c
A
E
D
a
a
b
b
a
a
a
c
a
a
a
c
a
b
c
c
Supplementary figure 7: Effect of melatonin priming on relative fold change expression of MAPK and necrosis related genes under optimal and PEG-induced stress conditions in drought sensitive (L-799) and drought tolerant (Suraj) varieties of cotton (Gossypium hirsutum L.). Relative fold change expression of necrosis related genes A) RAF18 B) BAG2, C) BAG6, D) MC9 and E) SOG1. The data represent mean values ± SD of 3 independent experiments, with 3 replicates per treatment in each experiment. Different alphabets represent significant differences among the treatments according to Duncan multiple range test (DMRT) at p-value ≤ 0.05.

### Slide 8
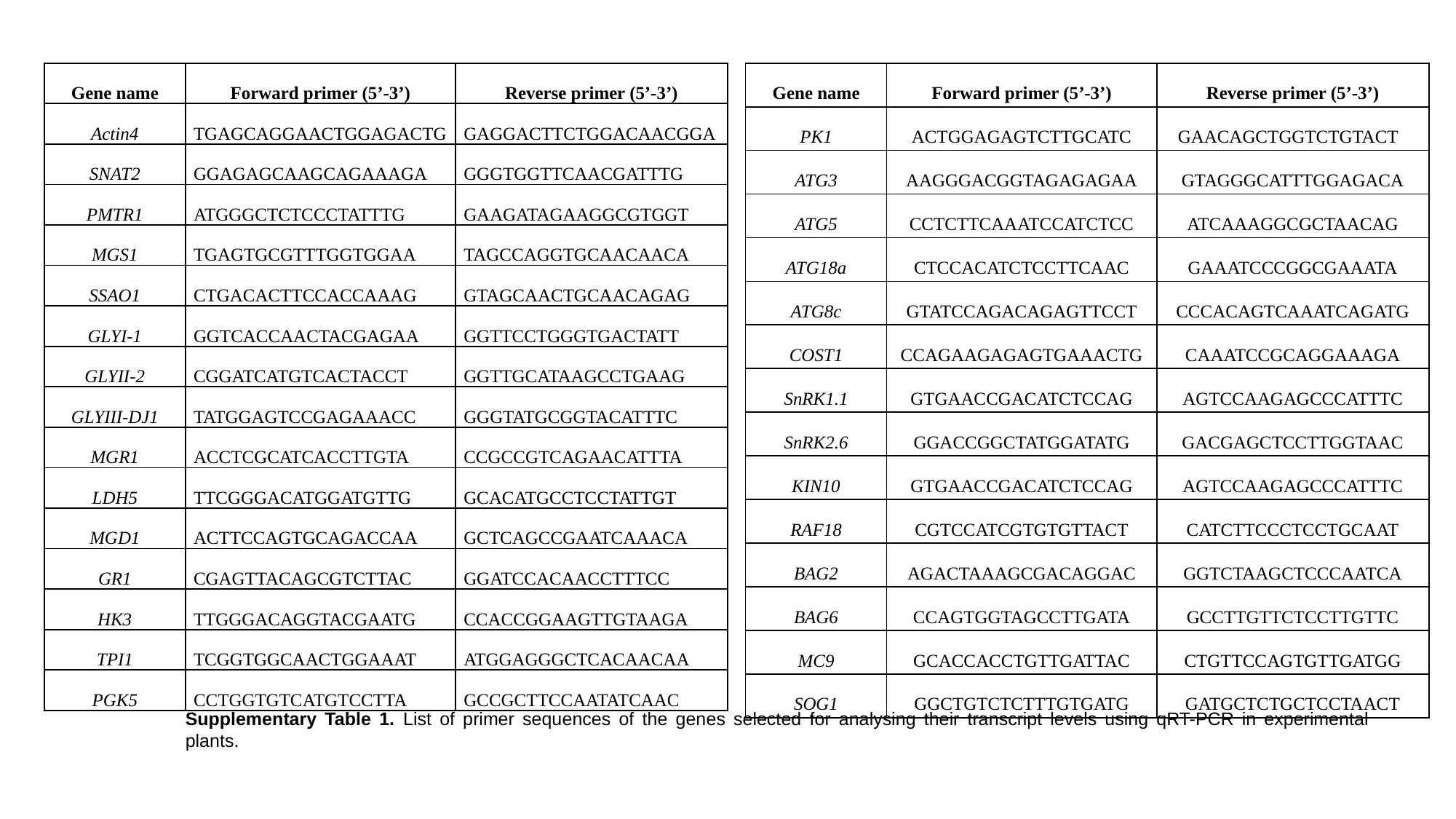

| Gene name | Forward primer (5’-3’) | Reverse primer (5’-3’) |
| --- | --- | --- |
| Actin4 | TGAGCAGGAACTGGAGACTG | GAGGACTTCTGGACAACGGA |
| SNAT2 | GGAGAGCAAGCAGAAAGA | GGGTGGTTCAACGATTTG |
| PMTR1 | ATGGGCTCTCCCTATTTG | GAAGATAGAAGGCGTGGT |
| MGS1 | TGAGTGCGTTTGGTGGAA | TAGCCAGGTGCAACAACA |
| SSAO1 | CTGACACTTCCACCAAAG | GTAGCAACTGCAACAGAG |
| GLYI-1 | GGTCACCAACTACGAGAA | GGTTCCTGGGTGACTATT |
| GLYII-2 | CGGATCATGTCACTACCT | GGTTGCATAAGCCTGAAG |
| GLYIII-DJ1 | TATGGAGTCCGAGAAACC | GGGTATGCGGTACATTTC |
| MGR1 | ACCTCGCATCACCTTGTA | CCGCCGTCAGAACATTTA |
| LDH5 | TTCGGGACATGGATGTTG | GCACATGCCTCCTATTGT |
| MGD1 | ACTTCCAGTGCAGACCAA | GCTCAGCCGAATCAAACA |
| GR1 | CGAGTTACAGCGTCTTAC | GGATCCACAACCTTTCC |
| HK3 | TTGGGACAGGTACGAATG | CCACCGGAAGTTGTAAGA |
| TPI1 | TCGGTGGCAACTGGAAAT | ATGGAGGGCTCACAACAA |
| PGK5 | CCTGGTGTCATGTCCTTA | GCCGCTTCCAATATCAAC |
| Gene name | Forward primer (5’-3’) | Reverse primer (5’-3’) |
| --- | --- | --- |
| PK1 | ACTGGAGAGTCTTGCATC | GAACAGCTGGTCTGTACT |
| ATG3 | AAGGGACGGTAGAGAGAA | GTAGGGCATTTGGAGACA |
| ATG5 | CCTCTTCAAATCCATCTCC | ATCAAAGGCGCTAACAG |
| ATG18a | CTCCACATCTCCTTCAAC | GAAATCCCGGCGAAATA |
| ATG8c | GTATCCAGACAGAGTTCCT | CCCACAGTCAAATCAGATG |
| COST1 | CCAGAAGAGAGTGAAACTG | CAAATCCGCAGGAAAGA |
| SnRK1.1 | GTGAACCGACATCTCCAG | AGTCCAAGAGCCCATTTC |
| SnRK2.6 | GGACCGGCTATGGATATG | GACGAGCTCCTTGGTAAC |
| KIN10 | GTGAACCGACATCTCCAG | AGTCCAAGAGCCCATTTC |
| RAF18 | CGTCCATCGTGTGTTACT | CATCTTCCCTCCTGCAAT |
| BAG2 | AGACTAAAGCGACAGGAC | GGTCTAAGCTCCCAATCA |
| BAG6 | CCAGTGGTAGCCTTGATA | GCCTTGTTCTCCTTGTTC |
| MC9 | GCACCACCTGTTGATTAC | CTGTTCCAGTGTTGATGG |
| SOG1 | GGCTGTCTCTTTGTGATG | GATGCTCTGCTCCTAACT |
Supplementary Table 1. List of primer sequences of the genes selected for analysing their transcript levels using qRT-PCR in experimental plants.
